# A humanized nanobody phage display library yields potent binders of SARS CoV-2 spike

**DOI:** 10.1101/2021.10.22.465476

**Authors:** Ying Fu, Juliana da Fonseca Rezende e Mello, Bryan D. Fleming, Alex Renn, Catherine Z. Chen, Xin Hu, Miao Xu, Kirill Gorshkov, Quinlin Hanson, Wei Zheng, Emily M. Lee, Lalith Perera, Robert Petrovich, Manisha Pradhan, Richard T. Eastman, Zina Itkin, Thomas Stanley, Allen Hsu, Venkata Dandey, William Gillette, Troy Taylor, Nitya Ramakrishnan, Shelley Perkins, Dominic Esposito, Eunkeu Oh, Kimihiro Susumu, Mason Wolak, Marc Ferrer, Matthew D. Hall, Mario J. Borgnia, Anton Simeonov

**Author notes:** These authors contributed equally: Ying Fu, Juliana da Fonseca Rezende e Mello, Bryan D Fleming and Alex Renn.

## Abstract

Neutralizing antibodies targeting the SARS-CoV-2 spike protein have shown a great preventative/therapeutic potential. Here, we report a rapid and efficient strategy for the development and design of SARS-CoV-2 neutralizing humanized nanobody constructs with sub-nanomolar affinities and nanomolar potencies. CryoEM-based structural analysis of the nanobodies in complex with spike revealed two distinct binding modes. The most potent nanobody, RBD-1-2G(NCATS-BL8125), tolerates the N501Y RBD mutation and remains capable of neutralizing the B.1.1.7 (Alpha) variant. Molecular dynamics simulations provide a structural basis for understanding the neutralization process of nanobodies exclusively focused on the spike-ACE2 interface with and without the N501Y mutation on RBD. A primary human airway air-lung interface (ALI) *ex vivo* model showed that RBD-1-2G-Fc antibody treatment was effective at reducing viral burden following WA1 and B.1.1.7 SARS-CoV-2 infections. Therefore, this presented strategy will serve as a tool to mitigate the threat of emerging SARS-CoV-2 variants.

## Introduction

The severe acute respiratory syndrome-coronavirus 2 (SARS-CoV-2) pandemic led to a worldwide emergency imposing massive strains on medical systems and a staggering number of deaths (1, 2). The scientific community responded with unprecedented celerity to develop effective vaccines conferring protective immunity (3). Although vaccination reduces the occurrence of severe disease and hospitalizations, there is potential for the emergence of escape variants and ongoing viral transmission (4). Thus, there remains an ongoing need for cost-effective, high-throughput, adaptable pipelines capable of identifying effective therapeutics against SARS-CoV-2 and other emerging pandemic threats.

SARS-CoV-2 entry into host cells relies upon the envelope-anchored spike (S) glycoprotein. This large trimeric class I protein densely decorates the viral surface, recognizes host angiotensin-converting enzyme 2 (ACE2), and contains the fusion machinery needed for viral entry (5, 6). During biogenesis, each S protomer in the mushroom shaped trimer is cleaved by cellular furin into S1 and S2 subunits responsible for target recognition and fusion, respectively. Each N-terminal S1 subunit contains a receptor binding domain (RBD) targeting the ACE2 on the host cell (7). The RBD is connected to a hinge that enables its transition between an ‘up’ state capable of binding to ACE2 and a ‘down’ state in which the interaction with the receptor is hindered by the proximity of the adjacent RBD. Receptor binding in the ‘up’ position triggers a cascade of events including the TMPRSS2-mediated cleavage of the stalk forming S2 protomers to reveal hydrophobic fusion peptides (FP) at each N-terminus of a long axial three helix bundle. Insertion of FPs into the membrane of the target cell is followed by a massive structural rearrangement resulting in its apposition with the viral envelope that leads to their fusion (8). Mutations in this targeting/fusion machine have been implicated in increased viral infectivity (9). As the driver of viral tropism, the RBD of the surface-exposed S protein is the focus of intense interest for the development of neutralizing antibodies and immunogens (10). Multiple variants of SARS-CoV-2 have arisen independently in which N501, one of several key contact residues within the RBD, has been mutated to tyrosine (N501Y). This mutation increases binding affinity to human ACE2 (11). These variants have multiple mutations in spike, which have been associated with increased transmission rates and reduced antibody neutralization (11). Structural insights into how antibodies bind to SARS-Cov-2 variants are still needed (12).

Anti-SARS-CoV-2 RBD neutralizing antibodies isolated from different platforms have been reported (13). Nanobodies are a unique class of heavy chain-only antibodies present in *Camelidae* and some shark species. Nanobodies (VHHs) and shark variable new antigen receptors (VNARs) are one order of magnitude smaller than their full-length IgG counterparts. The advantages of using nanobodies as anti-viral biologics are convenient bulk production at multi-kilogram scale in prokaryotic systems, long shelf-life, and greater permeability in tissues (14, 15). Nanobodies could be administered orally (e.g. V565 for gastrointestinal tract delivery) or through inhalation (e.g. ALX-0171), which are routes of administration that would be relevant for the COVID-19 pandemic (16, 17). The limitation of low affinity from synthetic nanobody libraires could be overcome by affinity maturation and/or constructing multi-valent/paratopic modalities (18).

Here, we report a rapid and efficient method of identifying neutralizing nanobodies directed against the SARS-CoV-2 from building nanobody phage display libraries, to *in vitro* and *ex vivo* high-throughput screenings. Synthetic humanized nanobody libraries were constructed by combining a framework with randomized complementarity-determining regions (CDRs) to diversify antigen recognition. Ten enriched RBD binders, among which the most potent, RBD-1-2G(NCATS-BL8125) and RBD-2-1F, exhibited an IC_50_ of 490 and 470 nM against SARS-CoV-2 pseudotyped particles, respectively. Constructing bi- or tri-valent formats of the RBD-1-2G domain resulted in improved affinity and neutralization potency against pseudotyped particles and authentic SARS-CoV-2 virus. Furthermore, cryo-electron microscopy (cryo-EM) structures for four nanobodies in complex with a stabilized SARS-CoV-2 S-protein ectodomain (19) revealed two distinct binding modes. The epitope for ‘Group 1’ nanobodies overlaps the receptor binding motif (RBM) at the distal end of the RBD. ‘Group 2’ binders targeted epitopes on the flat area proximal to the N-terminal domain of the adjacent monomer. This binding site was found to not overlap with the RBM and failed to inhibit ACE binding. Additionally, cryo-EM showed that three copies of RBD-1-2G were capable of binding to the same spike trimer. The RBD-1-2G nanobody was capable of neutralizing pseudotyped particles containing the N501Y mutation. A primary human airway air-liquid interface tissue model demonstrated that RBD-1-2G-Fc treatment was able to reduce viral burden in both WA1 and B.1.1.7 infections. Molecular dynamics simulation provides insights for nanobodies binding epitope exclusively focused on the spike-ACE2 interface. RBD-1-2G’s residues in the complementarity-determining region 3 (CDR3) contribute substantially to the neutralization of the wild-type (WT) SARS-CoV-2 and the B.1.1.7 alpha variant (N501Y). This approach could greatly apply to mitigate other infectious diseases.

## Materials and methods

### Cell lines

Cell lines used in this study were obtained from ATCC (HEK293 cells). ACE2-GFP HEK293T (CB-97100-203) cells were purchased from Codex Biosolutions. Expi293F cells with stable expression of human ACE2 (HEK293-ACE2) were custom produced by Codex Biosolutions (Gaithersburg, MD) (20). All cell lines used in this study were routinely tested for mycoplasma and found to be mycoplasma-free.

### Humanized nanobody library constructions

The DNA library of nanobodies was constructed by three-step overlap-extension PCR (OE-PCR). CDR mutagenesis were either designed based on analysis of nanobody and 892 human heavy chain of choice of amino acids in CDR3 from PDB since 2019 or IDT trimer 19 mix (21). Codon balanced libraries as shown in Figure 1A were synthesized by Glen Research and Keck Biotechnology Resource Laboratory. Primers were used to amplify the backbone of a humanized nanobody backbone of caplacizumab to generate a library of nanobody using vector pComb3XLambda (Addgene, Plasmid #63892). Following ligation, the DNA library was transformed into TG1 electrocompetent cells (Lucigen) with an efficiency of approximately 10^10^ CFU per library before amplification. To amplify the library, the phagemid library was added into 1 L 2TYAG medium (2% glucose, 100 µg/ml ampicillin) and shake at 30°C overnight. The overnight culture of library phagemids were collected through centrifugation at 3,500 rpm for 45 min. It was resuspended in 10 ml 50% 2TY/50% glycerol (v/v) to freeze down aliquots for making phage library. Each aliquot of 1.5 ml phagemids glycerol stock contains approximately 5x cell number of the library size. One aliquot is shaken at 37°C, 250 rpm until OD600 reaches to 0.9-1.0 in 1 L of 2TYAG medium, then 250 µl of M13KO7 helper phage (5×10^12^/ml) was added to each 1 L culture solution for a final concentration of 5×10^9^ and infect cells at 37°C for 60 min. The culture was centrifuged at 3500 rpm for 10 minutes the pellet was resuspended in 1 L 2TYKA (50 µg/ml kanamycin, 100 µg/ml ampicillin). The culture was shaken overnight at 30°C to produce phage particles. The supernatant containing the phage library was precipitated in 30% volume of PEG/NaCl overnight refrigerator and dissolved in total 12 ml PBS to reach a concentration at 10^14^/ml in PBS/glycerol stock (UV-Vis). To recapitulate different lengths of CDRs observed in nanobody VHH domains and human antibody heavy chains, different libraries containing different length of CDRs were made individually.

**Figure 1:**
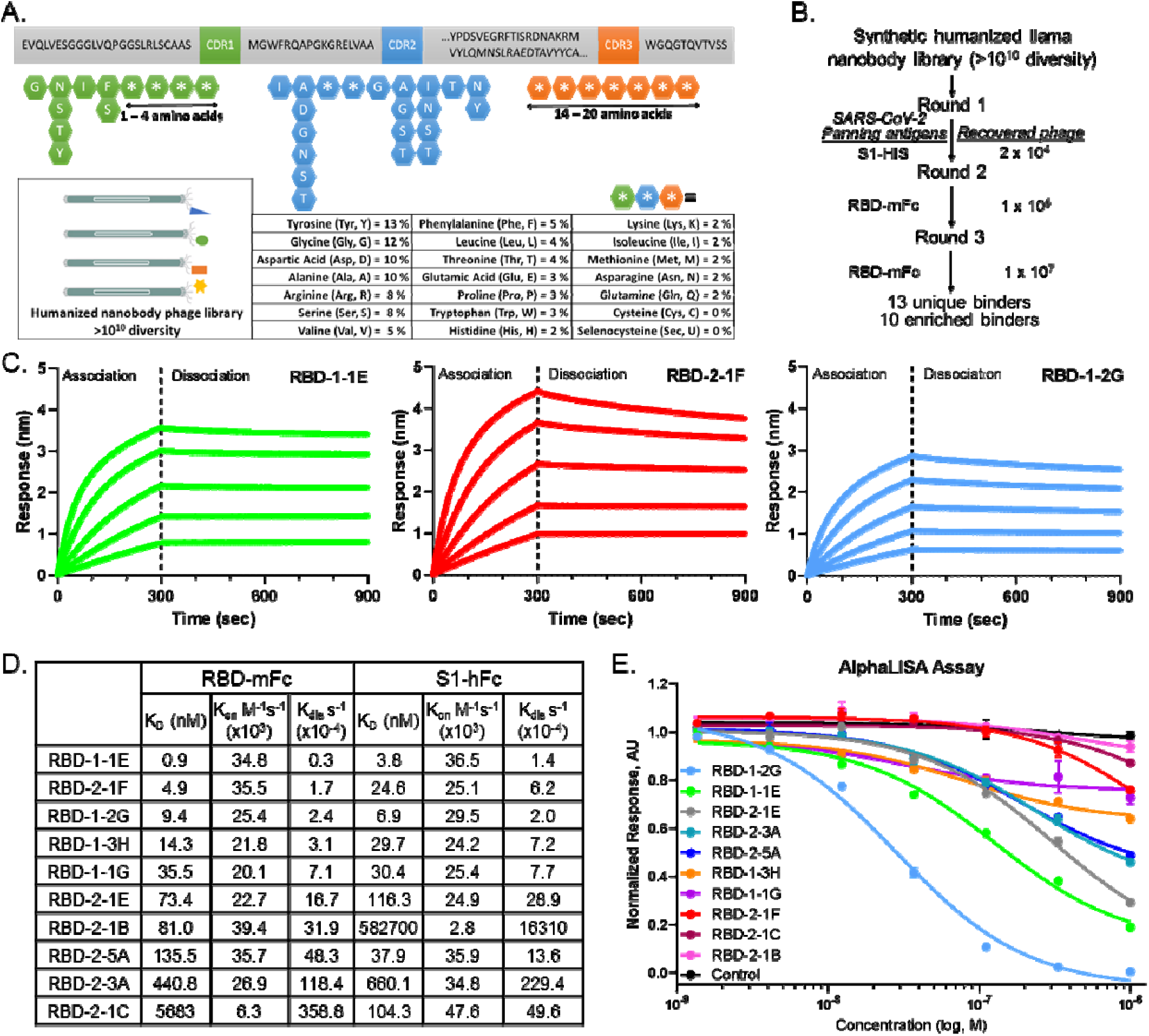
Discovery of anti-SARS-CoV-2 Spike RBD nanobodies that block interactions with ACE2. (A) Construction of a synthetic humanized llama nanobody library. (B) Selection strategy for identification of anti-RBD nanobodies using phage panning. (C) Bio-layer interferometry binding profiles of RBD-1-1E, RBD-2-1F and RBD-1-2G against RBD-mFc (200 nM to 12.5 nM, 1:2 dilution). (D) Association (k_on_) and dissociation (k_off_) rate constants and equilibrium dissociation constant (K_D_) of nanobodies binding to RBD-mFc and S1-hFc. Global fit calculations for RBD-1-1E, RBD-2-1F, and RBD-1-2G used (200 nM to 12.5 nM), with all others using 200 nM to 50 nM. (E) Nanobodies inhibition of RBD-Fc binding to ACE2-Avi using AlphaLISA.

### Isolation of RBD binders from phage library

On Day 1: 500 µl of 10 µg/ml S1-hFc or RBD-mFc was coated in a 5 ml immunotube (NUNC, 444202) overnight at 4°C. Day 2: A series of 10 libraries were mixed and used for phage panning (250 µL in EP tube). The libraries and the coated immunotube (pre-washed three times with PBS) were blocked with 10% (w/v) skim milk for 1.5 hours at RT. The immunotube was rinsed with PBS three times, then 0.5 ml pre-blocked phage solution was added before shaking at RT (300 rpm) for 1.5 hour. The immunotube was rinsed with PBST ten times, then PBS ten times to remove the unbound phage. Finally, RBD-bounding phage was eluted by incubating with 0.5 ml freshly made 100 mM triethylamine at RT for 15 mins. The eluted phage was further neutralized by 250 µl 1 M Tris-HCL buffer (pH 7.4). The neutralized phage (375 µL) was added to 3 mL TG1 (OD= 0.6 – 0.8) to infect at 37°C shaking for 60 min. The infected TG1 cells were collected at 3500 rpm for 10 min and spread in vented QTrays bioassay trays and titrations in small plates 2XYT agar plates with 2% glucose, 100 µg/ml ampicillin (Teknova, Y4295 and Y4204) incubate overnight at 30°C. On Day 3, All the colonies were scraped off the plates with 5 ml 2TY media and take 200 µL to grown in 25 mL 2TYAG until an OD600 of 0.5 – 0.8 is reached. At this point, helper phage (final concentration of 5×10^9^ per ml) was added and incubated with shaking at 37°C for 60 mins before changing the medium to 2TYKA for phage production at 30°C shaking overnight.

### Antibody and spike protein expression and purification

Nanobody-Fc format were produced through CRO company using HEK293 transient expression system (Sino Biological). Spike protein was expressed and purified using the method published previously (22). Nanobodies were expressed in *Vibrio natriegens* in auto-induction media, ZYM-20052, as outlined in Taylor *et al*. (23), with minor modifications for *Vibrio*. Specifically, media was amended with 1.5% (w/v) NaCl or Instant Ocean (Aquarium Systems), no lactose was added, IPTG induction began at an OD_600_ of 4 – 5, induction temperature was 30°C, and cells were harvested ∼6 – 8 hr after induction and frozen at −80°C. Frozen cell pellets were thawed and resuspended in 10 ml of PBS per 1000 optical density (OD_600_) units. Homogenized cells were lysed by passing thrice through a Microfluidizer at 9,000 psi. Lysates were clarified by centrifugation at 7,900 x g for 30 min at 4°C. Clarified lysates were filtered through 0.45 μM Whatman PES syringe filters. Nanobodies were purified using NGC medium-pressure chromatography systems (BioRad, Inc.). Clarified lysates were thawed, adjusted to 35 mM imidazole, and loaded at 3 ml/min onto HiTrap HP IMAC columns equilibrated in IMAC equilibration buffer (EB) of PBS, pH 7.4, 35 mM imidazole, and 1:1000 protease inhibitor cocktail. The columns were washed to baseline with EB and proteins eluted with a 20 column-volume (CV) gradient from 35 mM to 500 mM imidazole in EB. Elution fractions were analyzed by SDS-PAGE and Coomassie-staining. Positive fractions were pooled and further purified by size exclusion chromatography (SEC) using appropriately sized columns packed with Superdex 75 resin and equilibrated with PBS. After analysis of fractions from the SEC by SDS-PAGE and Coomassie-staining, appropriate fractions were pooled, concentrated in 10K MWCO Amicon Ultra centrifugation units, and snap frozen in liquid nitrogen (23).

### Nanobody trimer expression and purification

Protein expression constructs for production of trimeric nanobodies in insect cells were generated by synthesis of optimized DNA templates (ATUM, Inc.). Trimeric nanobody proteins were preceded by a honeybee mellitin (HBM) leader sequence for secretion and followed by a C-terminal His6 tag. DNA was flanked by attB1 and attB2 Gateway recombinational cloning sites and optimized for insect cell expression using ATUM’s algorithms. Templates were recombined using Gateway BP recombination (ThermoFisher) to generate entry clones, and subcloned into pDest-8, a pFastbac style baculovirus expression vector which utilizes the polyhedrin promoter to generate recombinant protein. Bacmid DNA was produced using the standard instructions for the Bac-to-Bac kit (ThermoFisher). Preparation of high-titer baculovirus stocks and protein expression protocols were essentially as described with minor changes (24). Specifically, the incubation temperature for the Tni-FNL cells, both before and after infection, was 27°C and the supernatants were collected after clarifying the cultures by centrifugation at 1700 x g for 15 min. The supernatant was then dialyzed against 1x PBS, pH 7.4 overnight at 4°C to remove any additives that might strip the nickel column. All purification processes were carried out using BioRad NGC Quest FPLC systems. The dialyzed supernatant was amended with 2 M imidazole to a final concentration of 25 mM imidazole and was then loaded at 5 ml/min onto a HiTrap HP IMAC column equilibrated in IMAC equilibration buffer (EB) of 1x PBS, pH 7.4 25 mM Imidazole. The column was washed to baseline with EB and proteins were eluted with a 20 column-volume (CV) gradient from 25 mM – 500 mM imidazole in EB. Elution fractions were analyzed by SDS-PAGE and Coomassie-staining. Positive fractions were pooled and further purified by size exclusion chromatography (SEC) using appropriately sized columns packed with Superdex S-75 resin and equilibrated with 1x PBS, pH 7.4. After analysis of fractions from the SEC by SDS-PAGE and Coomassie-staining, appropriate fractions were pooled, concentrated in a 10K MWCO Amicon Ultra centrifugation unit and snap frozen in liquid nitrogen.

### Affinity determination using bio-layer interferometry (BLI)

Affinity determination of VHHs were carried out using RBD-mFc or S1-hFc recombinant proteins (Sino Biological, 40592-V05H and 40591-V02H, respectively) as analytes. VHHs were loaded at 10 µg/ml in kinetics buffer (PBS with 0.1% protease-free BSA and 0.02% Tween-20 in PBS) onto Ni-NTA biosensors (Molecular Devices, ForteBio). Biosensors were hydrated for 10 minutes in water, then the plate was preincubated for 10 min at 30°C before the experiment started. Experimental parameters were baseline 1 min, loading 5 min, baseline2 3 min, association 5 min, dissociation 10 min. Association of RBD-mFc or S1-hFc was performed at 200, 100, 50, and 0 nM for all conditions, with 25 and 12.5 nM also included for the RBD-1-2G, RBD-2-1F and RBD-1-1E nanobodies. For data analysis, the 0 nM analyte with nanobody loaded was subtracted from corresponding nanobody loaded sensors. Alignment of curves to the last 5 seconds of baseline 2, inter-step correction aligned to dissociation and Stavitzky-Golay filtering was used. Data was processed, then a 1:2 bivalent analyte model with Global Fit was used for affinity calculations. (Data Analysis HT 11.1).

### Affinity determination RBD-1-2G multimeric modalities (BLI)

Affinity of bi- and tri-valent modalities of RBD-1-2G was determined against RBD-His (SinoBiological, 40592-V08H) using ForteBio amine reactive 2^nd^ generation (AR2G) biosensors. Biosensors were pre-hydrated in UltraPure DNase/RNase-free distilled water (Invitrogen, 10977015) for 10 minutes at 30°C before use. A baselined in water for 60 sec, followed by a 3 minute activation in a mixture of 20 mM 1-Ethyl-3-[3-dimethylaminopropyl] carbodiimide hydrochloride (EDC) and 10 mM N-hydroxysulfosuccinimide (s-NHS). Biosensor tips were then subject to 2.5 µg/ml of RBD-His in kinetics buffer (PBS with 0.02% Tween20 and 0.1% BSA) for 150 seconds or until an average binding of ∼1.5 nm was reached. Sensors were quenched using 1 M Ethanolamine (pH 8.5) for 150 sec. Another baseline of 180 sec in kinetics buffer was performed, before continuing to an association phase of 150 sec. Analytes RBD-1-2G, RBD-1-2G-Fc and RBD-1-2G-Tri, were prepared at 100 nM in kinetics buffer, then diluted 1:2 until 6.125 nM concentration was reached. Disassociation was measured in kinetics buffer for 300 sec. The 0 nM control wells were subtracted during data analysis. Data processing (Data Analysis HT 11.1) included alignment of curves to baseline2 (last 5 seconds), inter-step correction aligned to dissociation, and Stavitzky-Golay Filtering. Data were analyzed using a 1:1 binding model, with global curve fitting used to calculate apparent affinities.

### SARS-CoV-2 mutant S1 binding (BLI)

Determination of RBD-1-2G-Fc ability to bind S1 mutant variants was determine by loading of mutant protein onto an anti-human IgG Capture (AHC) biosensor, then exposing them to various S1 modalities at 200 nM. RBD-1-2G-Fc was prepared at 10 µg/ml in kinetics buffer (PBS pH 7.4 + 0.02% tween and 0.1% protease-free BSA). Wildtype S1 (Sino Biological, 40591-V08H) and B.1.1.7 variant (Sino Biological, 40591-V08H12) were prepared at 200 nM in in kinetic buffer (PBS pH 7.4 + 0.02% tween and 0.1% protease-free BSA) or 10 mM Acetate buffers with 150 mM NaCl (pH 4 – pH 6.0). Biosensors were hydrated for 10 min in water, and the plate was preincubated for 10 minutes at 30°C before the experiment started. Experimental parameters were baseline 1 min, conditioning 20 sec in 10 mM Glycine (pH 1.5), neutralization 20 secs, (condition and neutralization performed 3 times each), loading 2 min, baseline 1 min, association 1.5 min, dissociation 3 min. For data analysis, the 0 nM analyte with S1 protein loaded was subtracted from corresponding loaded sensors. Alignment of curves to the last 5 seconds of baseline 2, inter-step correction aligned to dissociation and Stavitzky-Golay filtering was used. Data were presented as the maximum response achieved during the association phase for the various pH conditions tested.

### AlphaLISA

The ability of nanobodies to disrupt recombinant spike protein RBD binding to ACE2 was assessed using an AlphaLISA (25). In dose response (1 µM – 6 pM) nanobodies were preincubated with 4 nM SARS-CoV-2 spike protein receptor binding domain fused to an Fc tag (RBD-Fc, SARS-CoV-2 spike protein residues 319-541) (Sino Biological, Wayne, PA) in PBS supplemented with 0.05 mg/mL BSA at 25°C for 30 minutes. After incubation ACE2 with a C-terminal His and AviTag (ACE2-Avi, human ACE2 residues 18-740) (ACROBiosystems, Newark, DE) was added to a final concentration of 4 nM and the resulting mixture was incubated at 25°C for 30 minutes. To produce the AlphaLISA signal a streptavidin donor bead, which recognizes ACE2-Avi, and Protein-A acceptor beads, which recognizes RBD-Fc, were added to a final concentration of 10 μg/ml each and the resulting mixture was incubated at 25°C for 40 minutes in the dark. AlphaLISA luminescent signal was measured using a PheraSTAR (BMG Labtech, Cary, NC) plate reader with a 384-well format focal lens equipped with an AlphaLISA optical module (BMG Labtech, Cary, NC). All experiments were performed in triplicate.

### QD_608_-RBD neutralization assay

QD_608_-RBD was synthesized as previously described (26). 25,000 ACE2-GFP cells were seeded into PDL-coated 96-well plates and incubated overnight. Nanobodies or nanobody-Fc constructs were pre-incubated with 10 nM QD_608_-RBD for 30 minutes. Cells were washed 1x with Optimem I Reduced Serum Media before 3 hour treatment with 10 nM QD_608_-RBD pre-incubated with nanobodies. Cells were imaged on the Perkin Elmer Opera using a 40x water immersion objective. 8-10 fields per well were captured from duplicate wells in a single plate. Approximately 1600 – 2500 cells were imaged per condition.

### Pseudotyped particle (PP) neutralization assay

SARS-CoV-2 pseudotyped particles (PPs) were purchased from Codex Biosolutions (Gaithersburg, MD), and were produced using a murine leukemia virus (MLV) pseudotyping system (27). All SARS-CoV2-S constructs was C-terminally truncated by 19 amino acids to reduce ER retention for pseudotyping. The WT S sequence was the Wuhan-Hu-1 sequence (BEI #NR-52420). The variant B1.1.7 (Alpha) contains mutations del69-70, del144, N501Y, A570D, D614G, P681H, T716I, S982A, and D1118H.

The PP neutralization assays were performed as follows. For WT PP assay, HEK293-ACE2 cells were seeded in white, solid bottom 1536-well microplates (Greiner BioOne) at 2000 cells/well in 2 µL/well media (DMEM, 10% FBS, 1x L-glutamine, 1x Pen/Strep, 1 µg/ml puromycin) and incubated at 37°C with 5% CO_2_ overnight (∼16 h). Test articles were titrated 1:2 in PBS and acoustically dispensed to assay plates at 200 nL/well. Cells were incubated with test articles for 3□h at 37°C with 5% CO_2_, before 2 µL/well of SARS-CoV-2-S PP was added. The plates were then spinoculated by centrifugation at 1500 rpm (453 xg) for 45 min, then incubated at 37°C for 48□h at 37°C with 5% CO_2_ to allow cell entry of PP and for the expression of luciferase reporter. After the incubation, the supernatant was removed with gentle centrifugation using a Blue Washer (BlueCat Bio). Then 4 µL/well of Bright-Glo Luciferase detection reagent (Promega) was added to assay plates and incubated for 5 min at room temperature. The luminescence signal was measured using a PHERAStar plate reader (BMG Labtech). For SARS-CoV-2 variants PP neutralization assays, HEK293-ACE2 cells were seeded in white, solid bottom 384-well microplates (Greiner BioOne) at 6000 cells/well in 15 µL/well media. The cells were incubated at 37°C with 5% CO2 overnight (∼16 h). Nanobodies were titrated 1:2 in PBS and added to cells at 1 µl/well. Cells were incubated with nanobodies for 1□hr at 37°C 5% CO2, before 15 µL/well of SARS-CoV-2 PP was added. The plates were then spinoculated by centrifugation at 1500 rpm (453 xg) for 45 min and incubated at 37°C for 48□hr at 37°C 5% CO2 to allow cell entry of PP and expression of luciferase reporter. After the incubation, the supernatant was removed with gentle centrifugation using a Blue Washer (BlueCat Bio). Then 20 µL/well of Bright-Glo Luciferase detection reagent (Promega) was added to assay plates and incubated for 5 min at room temperature. The luminescence signal was measured using a PHERAStar plate reader (BMG Labtech). Data was normalized with wells containing SARS-CoV-2 PP as 100%, and wells without PP as 0% entry. An ATP content cytotoxicity assay was performed by omitting the PP and adding media instead. Data was normalized with wells containing cells as 100%, and wells containing media only as 0%.

### SARS-CoV-2 cytopathic effect (CPE) assay

The SARS-CoV-2 cytopathic effect (CPE) assay was conducted in the BSL-3 facilities at the contract research organization (CRO) Southern Research (Birmingham, AL), as previously described (28). Briefly, nanobodies were titrated in PBS and added to 384-well assay plates at 1 µL/well to make assay-ready plates (ARPs), which are then frozen and shipped to the testing facility. Vero E6 African green monkey kidney epithelial cells (selected for high ACE2 expression) were dispensed into ARPs at 4000 cells/well in 25 µl of media (MEM, 1% Pen/Strep/GlutaMax, 1% HEPES, 2% HI FBS). The assay plates were incubated for 30 min, then inoculated with SARS-CoV-2 (USA_WA1/2020) at a multiplicity of infection (MOI) of 0.002 in media, and quickly dispensed into assay plates as 25 µL/well. The final cell density was 4000 cells/well. Assay plates were incubated for 72 h at 37°C, 5% CO2, and 90% humidity. CellTiter-Glo (30 µL/well, Promega #G7573) was dispensed into the assay plates. Plates were incubated for 10 min at room temperature. Luminescence signal was measured on Perkin Elmer Envision or BMG CLARIOstar plate readers. Data was normalized with buffer-only wells as 0% CPE rescue, and no-virus control wells as 100% CPE rescue. An ATP content cytotoxicity counter-assay was conducted using the same protocol as the CPE assay, without the addition of SARS-CoV-2 virus. Data was normalized with buffer-only wells as 100% viability, and cells treated with hyamine (benzethonium chloride) control compound as 0% viability.

### Membrane protein array assay

Membrane Proteome Array (MPA) screening was conducted at Integral Molecular (Philadelphia, PA). The MPA is a protein library composed of 6,000 human membrane protein clones, each overexpressed in live cells from expression plasmids. Each clone was individually transfected in separate wells of a 384-well plate followed by a 36 hr incubation (29). Cells expressing each individual MPA protein clone were arrayed in duplicate in a matrix format for high-throughput screening. Before screening on the MPA, the test antibody (RBD-1-2G) concentration for screening was determined on cells expressing positive (membrane-tethered Protein A) and negative (mock-transfected) binding controls, followed by detection by flow cytometry using a fluorescently-labeled secondary antibody. Each test antibody was added to the MPA at the predetermined concentration, and binding across the protein library was measured on an Intellicyt iQue using a fluorescently-labeled secondary antibody. Each array plate contains both positive (Fc-binding) and negative (empty vector) controls to ensure plate-to-plate reproducibility. Test antibody interactions with any off targets identified by MPA screening were confirmed in a second flow cytometry experiment using serial dilutions of the test antibody, and the target identity was re-verified by sequencing.

### Human Airway ALI model with qRT-PCR detection of viral mRNA levels

Day 20 normal human airway tracheobronchial (“epiairway”) tissue was purchased from MatTek Corporation (Ashland, MA) and cultured for 8 days in ALI, 37C, 5% CO2, to further mature the tissues in order to maximize for ciliated cell populations. Cells were grown at 37°C, with 5% CO_2_ levels in hangtop plates with 5ml Epiairway (MatTek, cat# AIR-100-MM) media in the basal chamber, with basal media changes every other day. On day 28, tissues were washed twice with 300ul of TEER buffer (MatTek) to remove excess mucus and dead cells prior to viral inoculation. The nanobody (10,000 nM), Fc (1,000 nM) or trimer (1,000 nM) were added directly to the tissues for five minutes prior to viral exposure. A total of 5,000 PFU of the WA1 or B.1.1.7 SARS-CoV-2 virus were added to the wells, followed by incubation at 37°C for 72 hours. Basal chamber media was replaced at 48 hours. 72 hours post infection, total RNA was isolated according to approved SOPs in the SARS-CoV-2 Virology Core BSL-3. Briefly, tissues were first dissociated into single cell suspensions by washing with 1ml of PBS for five minutes, followed by 5 minutes was in 200ul EDTA, followed by 200ul 0.25% trypsin EDTA for 10-15 minutes at 37C. Tissues were then broken into single cell suspensions by pipetting. 750 µl of Trizol LS was used to collect single cell suspensions per well and pipetted to homogenize the solution, followed by a ten-minute incubation at room temperature. RNA was purified using a Direct-zol RNA miniprep kit (Zymo Research) and RNA levels were determined by spectrophotometry.

The qRT-PCR reaction was conducted using the iTaq Universeral Probes one-step kit (Bio-Rad) supplemented with SybrGreen I nucleic acid stain (Invitrogen). Samples were prepared with 50 ng RNA per 10 µl reaction for the SARS-N reaction and 25 ng per well for the 18S reactions. Samples were analyzed on a Roche LightCycler 480 ii using Light Cycler 384 multi well plates. The following PCR protocol was utilized: reverse transcriptase step, 50°C for 10 minutes; Pre-Incubation step, 95°C for 5 minutes; Amplification step (45x), 95°C for 10 seconds, 60°C for 10 seconds, 72°C for 10 seconds; Melting curve step, 95°C for 10 seconds, 65°C for 1 minute, warm to 97°C with continuous fluorescent reading; cooling, 40°C for 30 minutes. The following primer pairs were used, SARS-N (Forward-TAA TCA GAC AAG GAA CTG ATT A -3’; Reverse-CGA AGG TGT GAC TTC CAT G -3’), 18S RNA (Forward-AAC CCG TTG AAC CCC ATT -3’; Reverse-CCA TCC AAT CGG TAG TAG CG -3’). Samples were run in technical triplicates that were averaged before data analysis. Samples were normalized to 18S mRNA loading to generate the delta Ct (dCt) values.

### SARS-CoV-2 prefusion S-protein ectodomain

The construct VRC 7471 was kindly provided by were generously provided by Dr. Kizzmekia Corbett and Dr. Barney Graham (Dale and Betty Bumpers Vaccine Research Center, NIAID). This construct encodes for nCoV S-2P-dFu-F-3C-H-2S,S ecto domain, with proline substitutions at residues 986 and 987, the Furin site mutated to GSAS, a T4 Fibritin trimerization motif, a PreScission protease cleavage site, and 8xHis and Strep tags. It is based on the one used for an early published structure (19). Expression was carried out in Expi293 cells (Thermo Fisher Scientific) following the manufacturer protocol. Briefly, 1 L Expi293 cells were transiently transfected using ExpiFectamine and incubated at 37°C for 18 hr. At this point enhancers were added and the temperature shifted to 32°C for 96 hr. To harvest the secreted protein, cells were pelleted by centrifugation at 400 g for 15 min and the supernatant was filtered through a 0.45 μm filter. A total of 5 ml of Talon Metal Affinity Resin (Takara Bio USA) was added to the supernatant and mixed overnight at 4°C. The supernatant/resin mixture was gravity loaded on a column and washed with 100 ml 50 mM Tris pH 8.0/150 mM NaCl followed by 100 ml of 50 mM Tris pH 8.0/500 mM NaCl. The Spike trimer was batch eluted with 50 mM Tris pH 8.0/150 mM NaCl/200 mM Imidazole. Fractions containing Spike trimer, as assessed by SDS-PAGE, were pooled and buffer exchanged/concentrated to ∼1 mg/ml using Amicon Ultra 4 centrifugal filters (Millipore Inc). Concentrated protein was divided into small aliquots, flash frozen in liquid nitrogen and stored at −80°C.

### Cryo-EM structure determination of the nanobodies bound to the trimeric spike protein

#### Specimen preparation

To increase hydrophilicity, grids were pretreated in a Tergeo EM Plasma Cleaner (PIE Scientific) under an argon atmosphere for 1 min, using the immersion mode at 25w. A 3 µL aliquot of 2.73 µM SARS CoV-2 S in Tris pH 8.0 50 mM and 160 mM NaCl buffer, mixed with each Nb was applied onto a clean UltrAufoil R1.2/1.3 300-mesh grid. Excess solution was removed by blotting with Whatman No. #1 prior to plunging into liquid ethane (−182°C) in a Leica EM GP 2 (Leica) vitrification robot (automated blot pressure, chamber at 25°C and 95% RH). The RBD-1-1G (1.5 mg/ml), RBD-1-2G (1.3 mg/ml), RBD-2-1F (1.8 mg/ml), RBD-1-3H (1.2 mg/ml), RBD-2-1B (1.7 mg/ml) and RBD-2-3A (4.2 mg/ml) were prepared in PBS (pH 7.4).

#### Data collection

Data was collected on either a Titan Krios or in a Talos Arctica transmission electron microscope (TFS) operated at 300 and 200 KeV respectively. The former equipped with a post column energy filter (Gatan) operated with a 20 eV slit size. Multiframe images were collected on direct electron detectors (Gatan K2). Details for each dataset are described in Supplemental Table 2.

#### Image processing

All image processing was carried out in the context of RELION-3.1.0. Motion correction was performed using the internal implementation and standard parameters without binning. CTF was estimated using CTFFIND4 in a resolution range of 3.0 – 30 Å, defocus range of 5,000 – 35,000 Å and a defocus step size of 500 Å. Laplacian-of-Gaussian detection with minimum and maximum diameters of 150 and 180 Å was used to pick particles. These were extracted using a box size of 300 pixels and Fourier down-sampled to 100 pixels (final size 3.562 A/pixel). Contamination and outliers were initially removed running 2 or 3 rounds of 2D classification. “Clean” particles were used to generate an initial 3D map using the stochastic gradient descent algorithm. Further elimination of outliers was achieved using 3D classification. A consensus C3 symmetric map was refined from each clean dataset and used for full defocus refinement (anisotropy, defocus and higher order aberration). A variety of approaches to classification (with/without alignment, symmetry expansion, etc.) were taken after this step in order to improve the resolution of the asymmetric portions of the spike.

#### Cryo-EM model building and analysis

An atomic model for the C3 symmetric closed structure of the SARS-CoV-2 spike protein (pdb: 6zp0) was used as the starting point for the fitting in the cryo-EM maps. To generate the “one-up” model, one of the RBDs in 6zp0 (residues 336 – 520) was manually aligned to the open state using pdb:6vyb as reference. The atomic model of the spike protein was positioned as a rigid body into each map using Fit in map tool on Chimera. After this, the atomic models were flexibly fitted to the maps using default real space refinement strategies (local grid search, global minimization, morphing and atomic displacement parameters refinement) on Phenix 1.18.2. This refinement was performed with Ramachandran restraints but without either secondary or NCS restraints.. The maximum number of iterations was increased to 150. The Fourier Shell Correlation plot between the atomic model simulated map and the cryo-EM map, were used to validate the fitting. In order to obtain the proper binding mode between each nanobody with the RBD, molecular docking calculations were combined with flexible fitting into the cryo-EM maps as follows. As the cryoEM map was not discernable enough for independent fitting of the nanobody to the RBD, 3D structural models of nanobody were generated using the I-TASSER program (30). The top 10 template structures of antibodies used by the threading program share 60 – 70% of sequence identities with the nanobody. All generated models showed a good quality in general (C-score > 0 and Z-score > 1). The best structural model was further refined with energy minimization and MD simulations using the Amber20 program (31). After fitting the spike atomic model to each density map, the Nb bound conformation of the RBD was extracted from the atomic model. The binding mode between nanobodies RBD-1-3H, RBD-2-1F, RBD-1-1G and RBD-1-2G to the corresponding fitted RBD was then predicted using ZDOCK server (See molecular dynamics below). All ten binding modes for each Nb-RBD complex reported by ZDOCK were fitted as rigid bodies to the corresponding RBD density maps in UCSF Chimera. An overview of the workflow can be found in Supplemental Figure S11.

#### Molecular dynamics

To better understand the interaction of RBD-1-2G with the RBM region, we performed atomic model fitting to identify the key binding residues. Since the resolution of the RBD region in our map was not high enough to directly derive an atomic model, fitting the atomic models of RBD and RBD-1-2G into the low-resolution map was the next best solution. However, the apparent symmetry of VHHs at low resolution leads to ambiguous assignments of their orientation in the context of the complex. A model of the spike (6zp0) was fitted into the map and the portion corresponding to the RBD coordinates was extracted (residues 328 to 531). Molecular docking calculations, where the RBD coordinates were kept rigid while the nanobodies’ were allowed to fluctuate were performed using the ZDOCK platform. Out of ten conformations requested, a pair that showed CDRs facing the RBM was positioned in the map relative to the RBD in the “half down” state of the spike model. The simulated density of the two complexes were compared with the map and the result exhibiting the highest correlation coefficient (CC) was selected for further refinement. This atomic model fitting strategy combined with molecular docking using the Cryo-EM map as a filter to choose the best predicted binding modes was tested against the nanobodies RBD-1-2G, RBD-2-1F, RBD-1-1G, and RBD-1-3H and showed successful predictions in all the cases (Data not shown).

Molecular dynamics simulations were performed on RBD-1-2G and WT RBD as well as RBD-1-2G and the alpha variant of RBD complexes to further characterize the energetics and the involvement of various residues located in the interacting interface. Starting protein structures were obtained from the cryo-EM density matched molecular docking. All MD simulations were performed using Amber.18 (12) with the ff14SB force field representing protein atoms. Initial minimization, equilibration, and all production runs were carried out with the GPU enhanced PMEMD module of Amber.18. Initial coordinates and topology files were generated using the Leap module, with the selection of rectangular TIP3P water box solvating the protein complexes. The box boundaries were extended at least 20 Å from the solute. Both systems contained 57 Na^+^ ions and 61 Cl^-^ ions that provided the charge neutralization and the 100 mM salt concentration. The WT system was comprised of 97,998 atoms and the mutant system had 98,006. After the initial equilibration of water and ions with each protein system was subjected to positional restraints with the 100 kcal/mol/Å^2^ force constant, a minimization was followed with the same force constants on the protein atoms. While maintaining the force constants at 10kcal/mol/Å^2^, each system was subjected to a 2 ns low temperature (T=100 K) constant pressure simulation to get the system density adjusted to a realistic value. After a step-wise slow heat-up to 300 K within 1ns, each system was further equilibrated for up to 10 ns with the position constraints on the protein atoms. Within the next 10 ns the positional constants were removed step-wise, and each system was subjected to a further 10 ns of equilibration. Three independent equilibrations were performed on each system (with the starting conformations for the second and third runs were selected from the 10th and 20th ns configurations of the first MD run). The constant pressure production runs were performed for 500 ns with the 2 fs time step for all systems. The particle mesh Ewald method was used in the treatment of long-range electrostatics with the short-range cut-off of 9 Å. The MMGBSA module of Amber.18 was implemented in free energy estimations with the selection of 0.15 M salt concentration and the default parameters (IGB = 5) within the Amber module. 500 configurations selected at each nanosecond of each trajectory were used in this calculation. All other analysis was done using the CPPTRAJ module of Amber.18.

## RESULTS

### Identification of SARS-CoV-2 neutralizing VHHs from synthetic, humanized nanobody libraries

To develop a platform for rapid nanobody discovery, we first designed a number of synthetic phage-displayed nanobody libraries, starting from a backbone derived from the first approved nanobody drug, caplacizumab (Fig. 1A). The caplacizumab framework region was combined with synthetized complementarity-determining loops (CDRs) to diversify the highly variable antigen-binding interface of the nanobody (21). This resulted in libraries each with diversity of >10^10^ transformants. Phage panning was performed for one round against S1 domain of SARS-CoV-2 spike, with two additional panning rounds against RBD of SARS-CoV-2 (Fig. 1B). A total of 13 nanobodies were identified during the panning process, with 10 of these showing sequence enrichment (Supplemental Table 1). Nanobodies were produced in *Vibrio natriegens* and purified using Ni-NTA affinity chromatography followed by size exclusion chromatography (Fig. S1A-B). Nanobody size and MW were confirmed by SDS-PAGE and high-resolution MS (Fig. S1C-D). bio-layer interferometry (BLI) analysis was used to visualize real time association and dissociation of RBD-mFc and S1-hFc to the various nanobodies. Profiles for RBD-1-1E, RBD-2-1F and RBD-1-2G can be found in Figure 1C. Global fit modeling revealed binding affinities of 0.9 nM (RBD-1-1E), 4.9 nM (RBD-2-1F) and 9.4 nM (RBD-1-2G) towards RBD-mFc (Fig. 1D, Fig. S2). When S1-hFc was used as the analyte, RBD-1-2G showed an affinity of 6.9 nM, a 26.6% improvement (6.9 nM vs 9.4 nM) over the RBD-mFc analyte. Reduced binding affinity was observed for both RBD-2-1F (24.6 nM vs 4.9 nM) and RBD-1-1E (3.8 nM to 0.9 nM) when switching to the S1 format (Fig. 1D, Fig. S3). The observed differences on affinity are probably due to the binding epitope on RBD being less accessible in the context of dimerized S1-hFc. To evaluate whether these nanobodies inhibit RBD-ACE2 complex formation, an AlphaLISA was developed (25). ACE2-Avi and RBD-Fc association was determined in the presence of increasing concentration of nanobodies to evaluate their receptor-blocking capabilities (Fig. 1E). Despite RBD-1-2G having the second highest affinity for S1-hFc, it yielded a lower IC_50_ than RBD-1-1E (28.3 nM vs. 126 nM) (Fig. 1E). These results support RBD-1-2G as a potential candidate for therapeutic development.

### Multimerization of RBD-1-2G to improve neutralization of SARS-CoV-2 virus

To further evaluate RBD-1-2G and to increase SARS-CoV-2 neutralization performance, we constructed bivalent and trivalent modalities. The bivalent format, RBD-1-2G-Fc, had an improved binding affinity of 1.9 nM compared to the monovalent format with 14.3 nM (Fig. 2A, Fig. 2B). The trivalent format (RBD-1-2G-Tri) was constructed through the linking of three monovalent formats with (GGGGS)_3_ flexible linkers. This further improved the apparent affinity to 0.1 nM for RBD binding (Fig. 2C).

**Figure 2:**
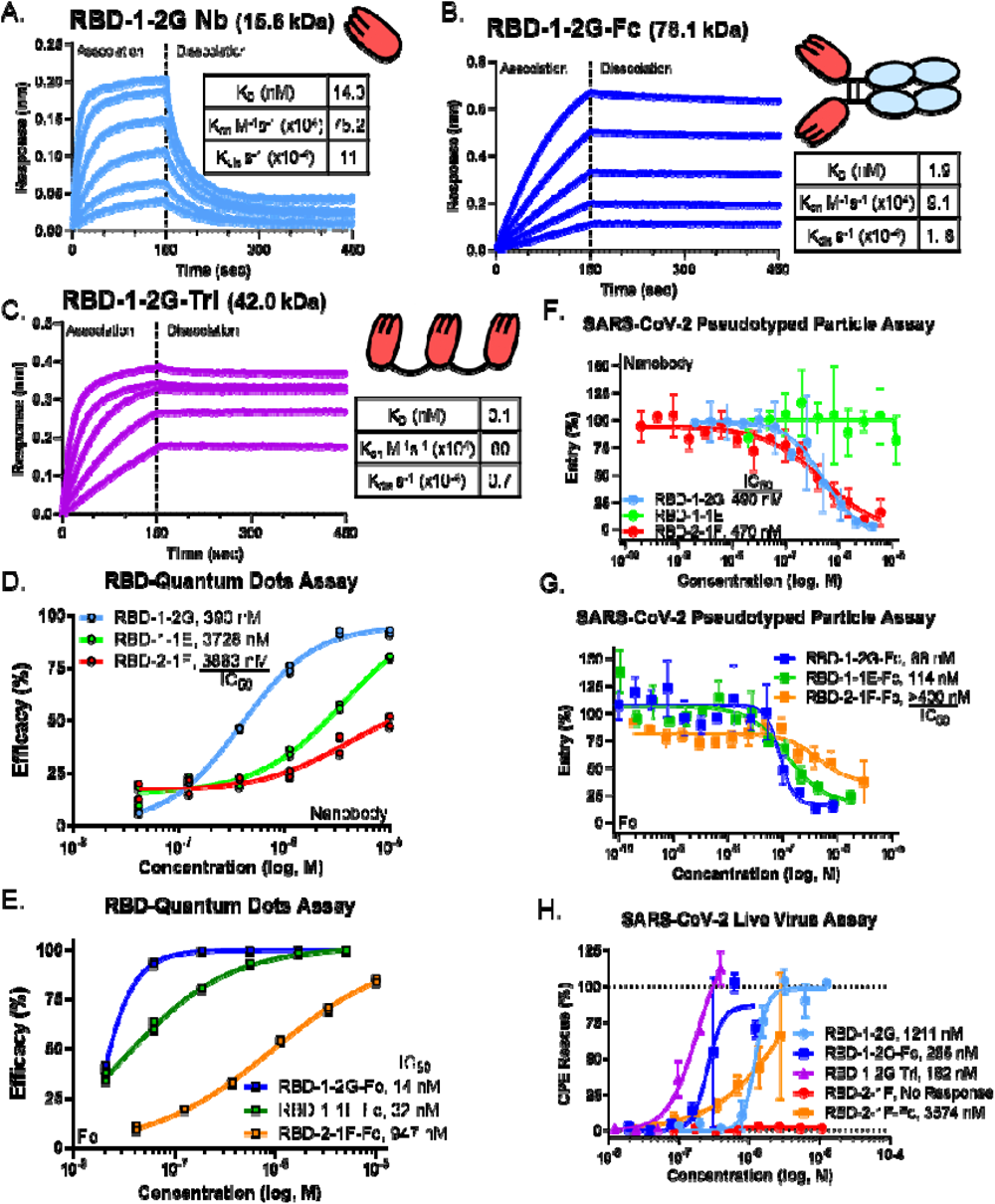
Multivalency improves affinity and inhibition of SARS-CoV-2 infection *in vitro*. (A-C) Bio-layer interferometry binding profiles for the (A) RBD-1-2G Nb, (B) RBD-1-2G-Fc and (C) RBD-1-2G-Trimer against RBD-His (100 nM to 6.25 nM, 1:2 dilution). (D-E) QD endocytosis assay using QD_608_-RBD and ACE2-GFP HEK293T cells to visualize receptor binding. Nanobody efficacy in reducing RBD internalization by (D) Nanobody and (E) F constructs. N = duplicate wells, approximately 2500 cells and 1600 cells, respectively. (F-G) SARS-CoV-2 pseudotyped particle entry assay using HEK293-ACE2 cells as target. Inhibition of pseudotyped particle entry was tested for Nanobody(F) and Fc(G) constructs Representative data from two independent experiments. Data represents mean inhibition per concentration (n=3), all error bars represent SEM. Inhibition of SARS-CoV-2 live virus infection with the RBD-1-2G and RBD-2-1F in various formats. Representative biological replicate with n = 2. Technical replicates are n = 2 per concentration, all error bars represent S.D.

To test if this improved affinity would translate to improved therapeutic potential, we first performed an endocytosis assay using SARS-CoV-2 RBD quantum dots (QD_608_-RBD) (Fig. S4, Fig. S5) (26). RBD-1-2G showed the highest potency with an IC_50_ of 390 nM (Fig. 2D). Conversion of RBD-1-2G to the Fc format reduced the IC_50_ to 14 nM (Fig. 2E). A similar trend was observed for RBD-1-1E-Fc (3728 nM to 32 nM), but not for RBD-2-1F-Fc (947 nM to 3883 nM) (Fig. 2D, Fig. 2E). Despite RBD-2-1F having a better affinity than RBD-1-2G as nanobodies (4.9 nM vs 9.4 nM, Fig. 1D), RBD-1-2G-Fc showed higher affinity than RBD-2-1F-Fc (Fig. S6). The binding of RBD-2-1F to RBD may be inhibited by the presence of a Fc domain.

*In vitro* neutralization assays using SARS-CoV-2 S-protein pseudotyped murine leukemia virus (MLV) vector particles were employed to further characterize the antiviral activity (27). RBD-1-2G was found to neutralize SARS-CoV-2 pseudotyped viruses with an IC_50_ of 490 nM (Fig. 2F). Significant improvements were seen with the Fc and trimer modalities showing inhibition at 88 nM (RBD-1-2G-Fc) and 4.1 nM (RBD-1-2G-Tri) (Fig. 2G, Fig. S7). Consistent with the QD internalization assay, RBD-2-1F-Fc showed improvement over RBD-2-1F but failed to neutralize better than the RBD-1-2G modalities (Fig. 2F). The other nanobodies and Fc modalities tested showed much less potency (Fig. S7). A SARS-CoV-2 live virus screen was performed to assess the neutralizing effect of nanobodies. RBD-1-2G-Tri produced an IC_50_ of 182 nM, followed closely by RBD-1-2G-Fc with an IC_50_ of 255 nM (Fig. 2H). The trimer and Fc modalities of RBD-1-2G were found to be more potent than RBD-1-2G (IC_50_=1211 nM) and the RBD-2-1F-Fc (IC_50_=3574 nM). These data support RBD-1-2G and its multimeric modalities for further development as SARS-CoV-2 therapeutics.

### RBD-1-2G displays low off-target binding

A human membrane proteome array (MPA, approx. 6,000 human membrane proteins + SARS-CoV-2 spike protein) was used to screen and evaluate potential poly-reactivity and non-specificity of RBD-1-2G. The MPA screen confirmed high-specificity binding of RBD-1-2G to SARS-CoV-2 spike with minimal binding to extraneous human membrane proteins (Fig. S8A). Binding to *mitochondrial elongation factor 1* (MIEF1), an intracellularly expressed protein, was detected in an initial screen. Validation of this target revealed that RBD-1-2G bound MIEF1-transfected cells in a similar fashion to vector-transfected cells (Fig. S8B). Additionally, since MIEF1 is a mitochondrial protein, it will be unlikely to interfere with a blood infusion or nebulized administration of RBD-1-2G. These assays suggest a minimal potential off-target effects of RBD-1-2G if selected for further development.

Additionally, we lyophilized RBD-1-2G nanobody and tested the reconstituted nanobody in a similar live virus study. The lyophilization process had little effect on overall activity in the live virus screen, with IC_50_ measuring 5.9 µM for the lyophilized and 14.8 µM in untreated samples (Fig. S9). The ability of RBD-1-2G to retain activity after lyophilization is a positive feature that can be useful in the manufacturing and formation of nanobody-based therapeutics.

### Cryo-EM revealed two distinct modes of binding for neutralizing nanobodies

To uncover the region bound by nanobodies, we used single particle cryo-EM to obtain electron scattering density maps of a soluble form of S-protein ectodomain in complex with 4 nanobodies (Fig. 3). Maps of the complexes with RBD-1-2G, RBD-2-1F, RBD-1-1G and RBD-1-3H featured extra density in the RBD region (Fig. 3A). Two additional nanobodies (RBD-2-3A and RBD-2-1B) were tested, but the observed electron densities were found to be similar to unliganded spike (Data Not Shown). Low affinity was also observed when these nanobodies bound S1-hFc by bio-layer interferometry (Fig. 1D), suggesting their epitopes are not accessible in the context of the trimeric spike.

**Figure 3:**
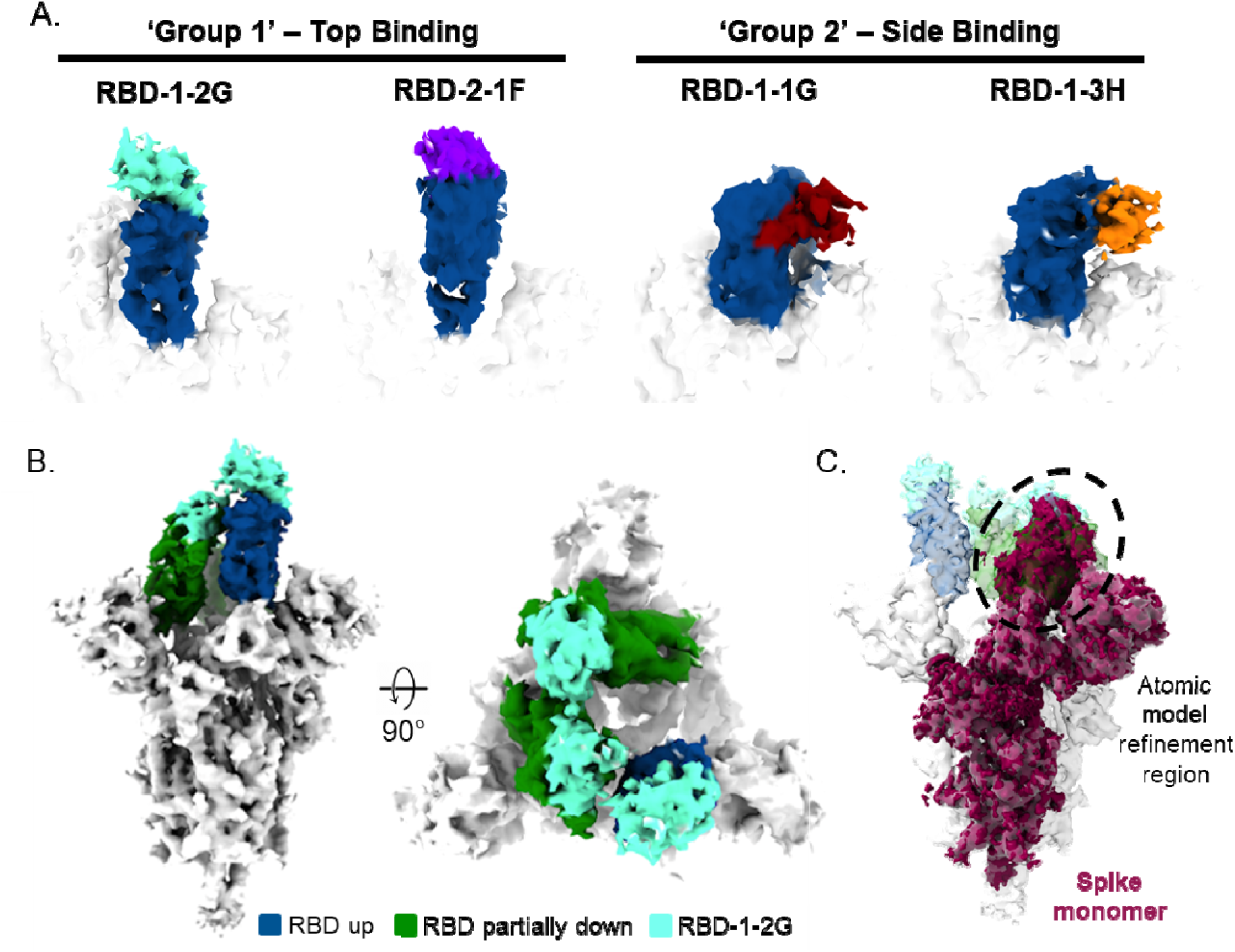
Cryo-EM of SARS-CoV-2 spike trimer and RBD binding nanobodies. (A) Cryo-EM analysis of nanobodies complexed with the RBD (up state) revealed two distinct binding modes. (B) Top and side views of the cryo-EM map of S-protein in complex with 3 molecules of RBD-1-2G (cyan). (C) Side view of the S-protein highlighting the spike monomer region refined during image processing is shown. The density containing the RBD in the laid state was used for the atomic model fitting refinement.

The accessible epitopes can be classified into two groups. ‘Group 1’ comprises the binding area for RBD-1-2G and RBD-2-1F located at the distal end of the ‘up’ RBD in an area that overlaps with the RBM. Nanobodies RBD-1-1G and RBD-1-3H bind to epitopes in ‘Group 2’, exposed on the external face of the erect RBD in an area not overlapping the RBM (Fig. 3A). The atomic model fitting of the spike (PDBID: 6zp0) suggest that the ‘Group 1’ binders overlap with the ACE2 binding site, while ‘Group 2’ binders do not prevent ACE2 binding (Fig. S10). In the down conformation, epitopes of ‘Group 2’ on the RBD are sterically occluded by the NTD of the neighboring monomer, thus nanobodies of this group were only detected when bound to the RBD in the ‘up’ conformation. In contrast, densities corresponding to RBD-1-2G can be detected in the ‘up’ and ‘partial down’ conformations of the RBD. This unique intermediate between the ‘up’ and ‘down’ states allows for the accommodation of three molecules of RBD-1-2G without steric hindrance (Fig. 3B), likely explaining the high activity for RBD-1-2G. In order to improve resolution of the binding area, independent refinement of “RBD down” spike monomers (shown in maroon in Fig. 3C) was carried out using symmetry expansion and signal subtraction with the spike monomer. The obtained map was used the atomic model fitting.

### RBD-1-2G binds to WT and B.1.1.7 variant (Alpha) at various physiological pHs

Recently, different strains of SARS-CoV-2 with N501Y mutation have been identified which have been associated with increased transmissibility and a reduction in neutralizing antibody activity (11, 32). We exposed biosensor-immobilized RBD-1-2G-Fc to both wildtype and B.1.1.7 mutant (N501Y) S1-His and evaluated the BLI signal. The maximum binding response to WT or B.1.1.7 S1-His during the association phase was graphed over a range of pH conditions (7.4 – 4.0) (Fig. 4A). Similar binding profiles were observed for WT and B.1.1.7, but a stronger signal was observed with B.1.1.7. Strongest bindings were observed between pH 5.0 and 6.0. Binding was detectable down to a pH of 4.5. A similar pattern was observed when RBD-His was used instead of S1-His, although there appeared to be stronger binding to the WT (Fig. 4B). The strongest binding was observed at pH 6.0 with a gradual reduction as pH was lower to 4.0. These data would suggest that the RBD-1-2G may stay bound to SARS-CoV-2 WT and B.1.1.7 variant (Alpha) after internalization while in the endosome (pH = 6.5 – 4.5).

**Figure 4:**
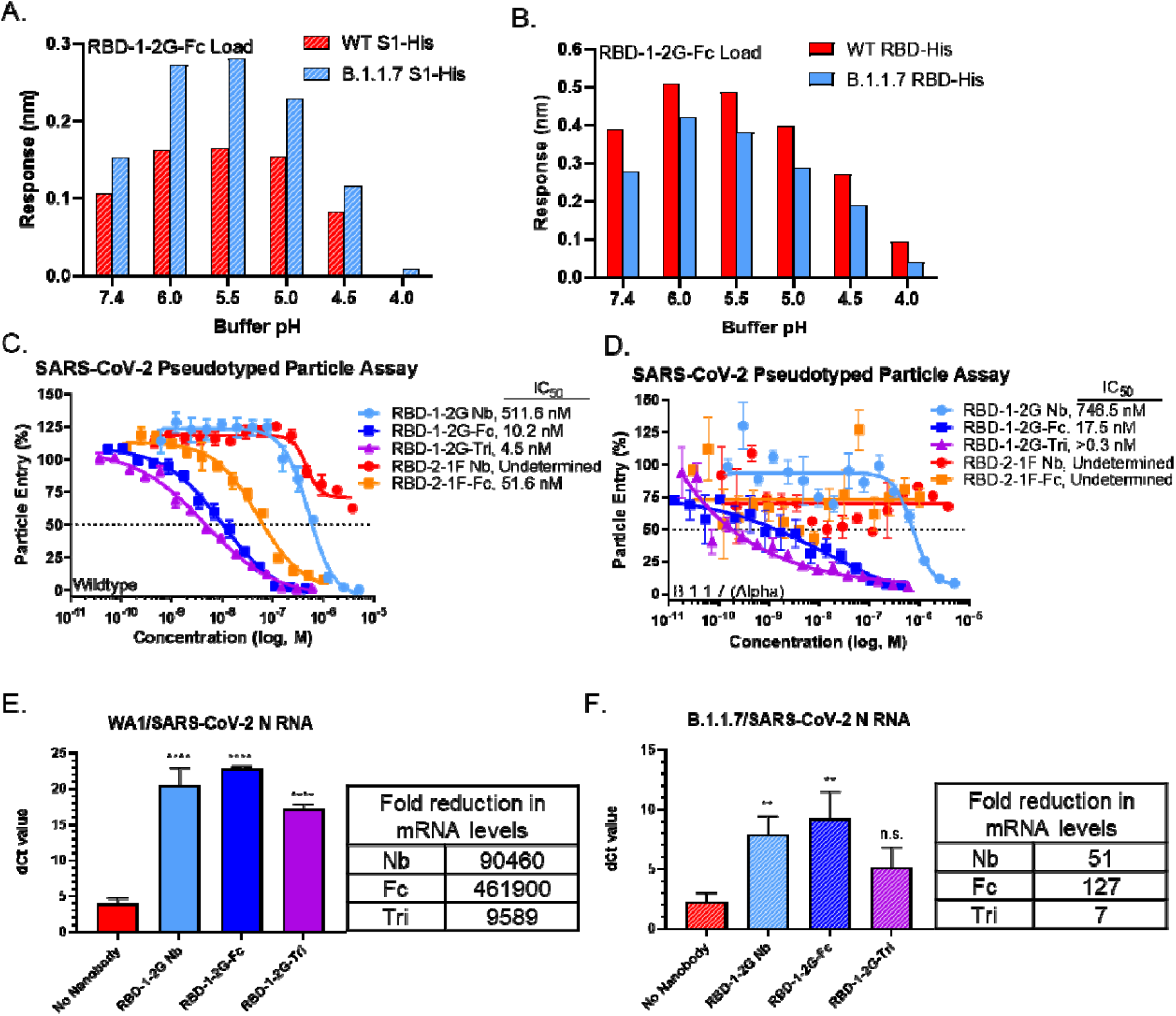
Binding and neutralization of RBD-1-2G to the WT and B.1.1.7 variant (N501Y). (A) Maximum response values reached during the association phase by RBD-1-2G-Fc binding wildtype (WT) and B.1.1.7 (Alpha) variant S1-His. Differences in pH were achieved using PBS (pH 7.4) or 10 mM Acetate buffers with 150 mM NaCl (pH 4 – pH 6.0), all buffers contained 0.1% BSA and 0.02% Tween. (B) Maximum response values reached during the association phase by RBD-1-2G-Fc binding WT and UK variant RBD-His proteins. (C-D) SARS-CoV-2 pseudotyped particle entry assay using HEK293-ACE2 cells as target. Inhibition of WT(C), B.1.1.7 (D) pseudotyped particle treated with various RBD-1-2G and RBD-2-1F formats. Representative biological replicate with n = 2. Technical replicates are n = 3 per concentration, all error bars represent S.D. (E-F) Primary human airway air liquid interface (ALI) model of SARS-CoV-2 infection. Treatment was added at 10,000 nM for nanobody treatment and 1,000 nM for Fc and trimer modalities. Levels of SARS-CoV-2 N mRNA following infection with WA1 (E) or B.1.1.7 (F) SARS-CoV-2 viral infection was determined by qRT-PCR and normalized to 18S mRNA levels. Bars represent that average dCt from biological triplicates, errors bars represent S.D. Fold reduction in mRNA levels are compared to the no nanobody infection control. One-way ANOVA was used to compared treatment groups with the no nanobody control. Significant *p* values are represented as follows: \*\**p* < 0.01, \*\*\*\**p* < 0.0001, n.s. *p* > 0.05.

To better judge how the RBD-1-2G formats would perform as therapeutic agents, we performed a pseudotyped particle assay to compare their neutralization ability against the wild type and B.1.1.7 spike proteins. RBD-1-2G-Tri was found to inhibit the B.1.1.7 pseudotyped particles better than the wildtype particles (0.3 nM vs 4.5 nM) (Fig. 4C, Fig. 4D). RBD-1-2G-Fc (17.5 nM vs 10.2 nM) and RBD-1-2G (746.5 nM vs 511.6 nM) showed similar IC_50_ against the B.1.1.7 variant (Fig. 4C, Fig. 4D). Additionally, RBD-2-1F-Fc was able to inhibit the WT pseudotyped particles but failed to inhibit the B.1.1.7 (Alpha) mutant version (Fig. 4C, Fig. 4D). To better replicate the complex tissues of the mammalian respiratory tract, we designed a 3D tissue model utilizing reconstructed normal human airway (tracheobronchial) air-liquid interface (ALI) tissues. We infected the human airway ALI tissues with 5,000 plaque forming units (PFU) for an approximate multiplicity of infection (MOI) of 0.005 of either WA1 or B.1.1.7 SARS-CoV-2 virus per well, in the presence or absence of the RBD-1-2G nanobody, Fc and trimer constructs. The monovalent nanobody was used at 10,000 nM, with the multivalent Fc and trimer being administered at 1,000 nM. After 72 hours of infection the total RNA was collected, and viral mRNA levels of SARS-N was determined by qRT-PCR. Comparison of the delta Ct values (dCt) revealed a significant reduction in viral mRNA present in the wells. On average the dCt of the WA1 infected cells was around 4 cycles in the virus only treatment group. This was increased to 20.4 (Nanobody), 22.8 (Fc) and 17.2 (Trimer) cycles following treatment (Figure 4E). A similar trend was observed with the B.1.1.7 infected cells, but the cells were found to have higher burdens of viral mRNA. The averaged dCT for the no treatment group was found to be 2.2 cycles, indicating that these cells had roughly 3.5 times the viral mRNA than the WA1 infected wells, indicating a higher infection in B.1.1.7 SARS-CoV-2 infected airway ALI tissues in comparison to WA1 SARS-CoV-2 infected tissues. Treatment of the B.1.1.7 infected cells increased the dCT to 5.7 (Nanobody), 7.0 (Fc) and 2.9 (Trimer) (Figure 4F). The RBD-1-2G-Fc showed better activity than the trimer in the ALI tissue model, indicating that the Fc format might be the most favorable for clinical transition.

### Atomic modeling of interactions between RBD-1-2G and RBM

To elucidate the residues that participate in the binding of RBD-1-2G with the RBD regions of WT and B.1.1.7 (Alpha) variant, molecular docking of the RBD-Nanobody(Nb) complex was employed before fitting it to the spike (Fig. S11). The best binding mode was selected by superimposing the docking results with the cryo-EM maps. Next, triplicates of the RBD-1-2G-WT RBD and RBD-1-2G-B.1.1.7 RBD complexes were subjected to over 1 µs molecular dynamic (MD) simulations in solution.

The root mean square deviations (RMSDs) of the complexes in all simulations were low, confirming that the systems were stable (Fig. S12). The consistent pattern of the curve, showing mainly a single jump, indicating at least two possible distributions in both systems (Fig. S12, SET2 and SET3). The highly similar patterns of interactions contribute to a very similar binding mode of RBD-1-2G to both the WT and B.1.1.7 (Alpha) variant. The MD results showed that RBD-1-2G binds in a similar angle to both the WT and B.1.1.7 (Alpha) variant (Figure 5A and 5B). When comparing the position fluctuations per residue in each complex, similar patterns in root mean square fluctuations (RMSFs) (Fig. S13) emerged in both systems, with larger variations shown in two distinct regions bracketing the RBD residues 355 – 375 and 470 – 490 (Fig. 5A, 5B, S13A, S13B).

**Figure 5:**
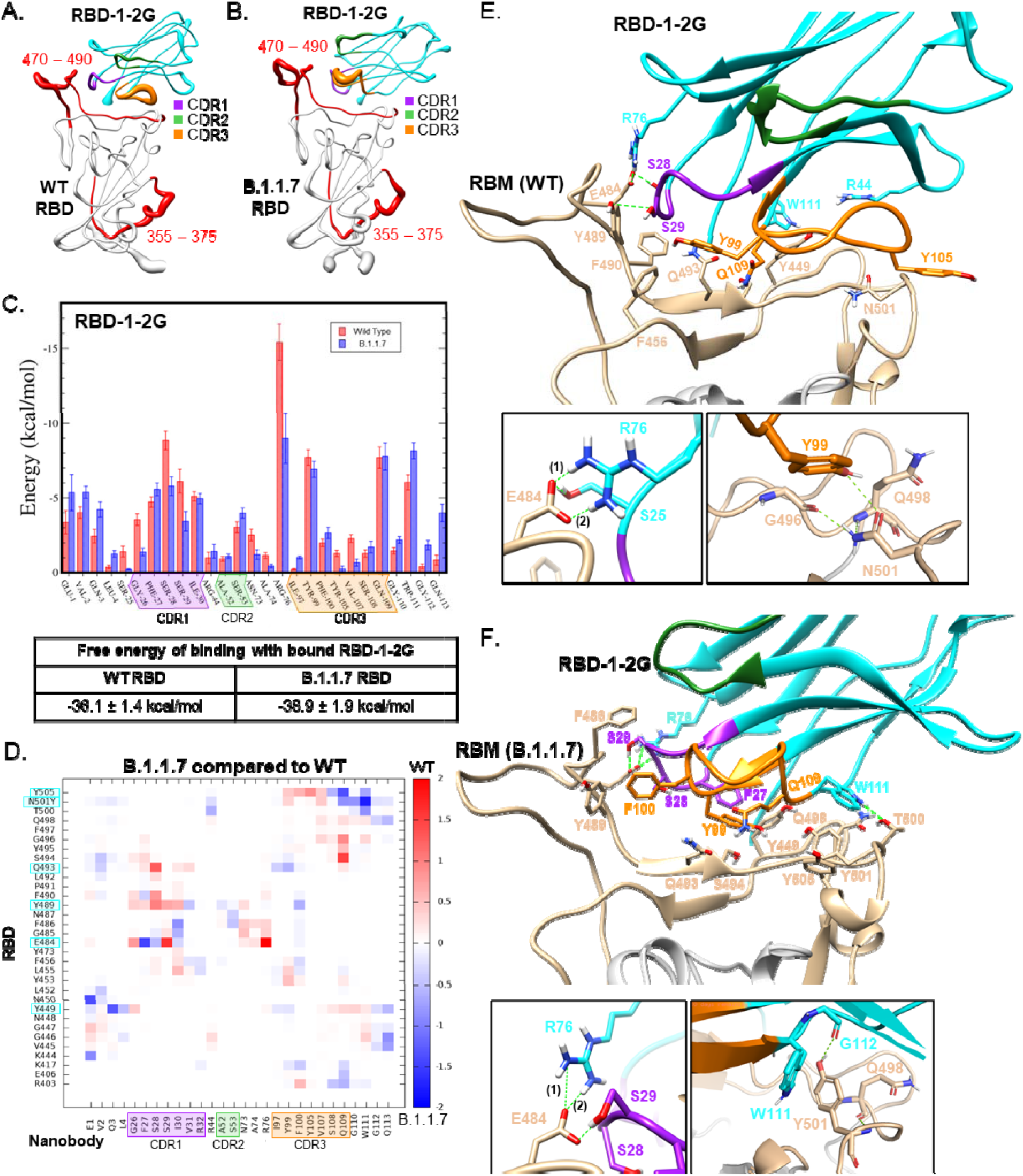
Molecular dynamics of RBD-1-2G with Wild type and B.1.1.7 RBD variant. (A-B) Sausage plot representation of RBD-1-2G in complex with (A)WT RBD and (B) B.1.1.7 RBD. The most flexible regions are indicated in red. (C) Free Energy of binding contribution of each Nb residue in complex with the RBD WT and B.1.17 variant. (D) Heatmap comparing RBD-1-2G’s free energy of binding contributions against the B.1.1.7 and WT RBD. RBD residues that contributed greater than -5 kcal/mol are highlighted with cyan. (E-F) Atomic model obtained after the atomic model fitting and MD simulations of RBD-1-2G in complex with (E) WT RBD and (F) B.1.1.7 RBD.

CDR1 and CDR3, as well as a few residues in the constant region (R76, N73, and W111), were mainly responsible for the stabilization of the complex based on the individual energies from the MD simulation (Fig. 5C). A few residues in the N-terminal constant region (E1, V2, Q3) and CDR2 (A52 and S53), interact with the RBD in both WT and mutant forms (Fig. 5C). The MM/GBSA method was used to estimate the binding free energy (ΔG binding) for nanobody– RBD complex systems. One thousand configurations were selected at 20 – 30 ns time intervals throughout the MD simulation trajectories to compute the MM/GBSA total free energy difference. As shown in Fig. 5C, the average calculated binding energies of RBD-1-2G in complex with WT RBD is -36.1 kcal/mol, while the B.1.1.7 RBD is −38.9□kcal/mol. A slight increase in the number of pairs of hydrogen bonds or salt bridges was observed in the complex with the mutant RBD compared with the WT complex (Fig. S14). Thus, RBD-1-2G can form stable complexes with both the WT and B.1.1.7 variant, with the variant being slightly favored. The predicted interaction energy provided by each residue showed that the salt bridge between E484 and R76 had the most significant contribution to the free energy of binding in both simulations (Fig. 5C, S15). F27 shows increased interaction with E484 in the B.1.1.7 variant, while S29 and R76 showed decreased interactions (Fig. 5D, S15). The observation of similar binding modes and the activity of these nanobodies against the WT and B.1.1.7 variant is conferred collectively by the above results. However, the conformation adopted for RBD-1-2G in the WT complex shows the salt bridge between E484 and R76 is further stabilized by the interaction of E484 with S25, S28, S29, and G26 in some instances (Fig. 5D, S13, Table S3). These stabilizing interactions were also observed with the B.1.1.7 variant in addition to N73 that was also found to play a role. Interestingly, the N501 residue found in the WT SARS-CoV-2 strain was found to interact with Y99 in the CDR3 region of RBD-1-2G (Fig. 5E). However, when the spike contains the N501Y mutation, the primary interaction residues switch to W111 and G112 to interact with the new tyrosine residue (Fig. 5F). This flexibility in the CDR3 region is characteristic of nanobody binding. This flexibility and ability to compensate for spike protein mutations makes the RBD-1-2G nanobody and the multivalent constructs good choices for therapeutic development.

## DISCUSSION

In this study, we report the isolation and characterization of SARS-CoV-2 neutralizing single-domain antibodies from a pool of humanized phage libraries. These nanobodies bind the SARS-CoV-2 spike RBD with single-digit nM to µM affinity, and are capable of neutralizing S-protein pseudotyped and authentic viruses in mammalian cell models of SARS-CoV-2 infection.

RBD-1-2G showed the best overall potency and improved viral neutralization when incorporated into bivalent and trivalent modalities. RBD-1-2G binds an epitope on the top of the RBD that overlapped with binding site for ACE2 (Fig. S10). The potency observed for RBD-1-2G was not only attributed to its high affinity, but also due to the ability of this nanobody to bind the RBD in a wide range of conformations from the “up” to the “down” states (Fig. 3B). Recent cryo-EM studies revealed two prevalent states of S trimer: three RBD domains in “down” conformation which may indicate a conformational immune escape mechanism of action (6), or only one RBD in the “up” conformation corresponding to the ACE2-accessible state (6, 33). Some nanobodies (Group 2) could not be imaged in multiple RBD conformations, which would suggest that conformation switching to a protected, three RBD down conformation may sterically restrict binding. The ability of RBD-1-2G to bind in both the “up” and “half down” states allow three nanobodies to bind to a single spike trimer regardless of the RBD conformation state.

The molecular dynamics results provided a deep understanding of how RBD-1-2G interacts with the RBM. This molecule displayed at least two different binding modes, according to the RMSD analysis, which can be related with the low resolution observed in the Cryo-EM map specifically in the RBD and Nb region (Fig. S12). This Nb showed a very similar binding mode to the WT and B.1.1.7 variant, which was also observed in the Cryo-EM maps (Fig. 3B and S16). One of the reasons for the higher affinity of RBD-1-2G to B.1.1.7 RBD could be related to the presence of the tyrosine in position 501, since it is a longer residue than the asparagine residue found in the WT RBD. The tyrosine also displays higher rotational degree of freedom than the asparagine where the sidechain may be slightly dipolar in nature. Even having a flexible side chain, a dipolar asparagine sidechain may not show the same adaptability of the possible hydrogen bonds donors/acceptors as tyrosine. The mutant might favor the formation not just of the hydrogen bond with the Gln109 (as seen in WT) but also with the neighboring residues Gly112 and Trp111, thus slightly increasing the affinity. Overall, the high activity profile and the possibility of binding in different conformations to the RBD, supports the versatility of RBD-1-2G as a good prototype to design new candidates that are capable to adapt to different SARS CoV-2 variants.

The continued spread of SARS-CoV-2 infections around the world, primarily in dense population settings, has resulted in the emergence of new mutant lineages. For this study we used the B.1.1.7/Alpha (N501Y), which enhanced the RBD binding to ACE2 receptor, to determine the effect of the mutation on our nanobody. This variant (in early June 2020) was the dominant strain in the United States (>70% of all sequenced strains). The N501Y mutation has been found in newer variants including the current most dominant strains (11). RBD-1-2G was shown to bind both to the N501Y mutant and the WT RBD and that the hydrogen bond interaction with residue 501 is conserved. Nanobody have been shown to be able to recognize epitopes often inaccessible to conventional antibodies which could offer some advantages to target variants (34). It has been more challenging to screen antibodies which can bind to RBM but tolerate a mutation. In this work, we demonstrated the mechanism by which a nanobody targeting RBM could accommodate the N501Y mutant. This could pave the road to target other current and future variants. The ability to rapidly screen artificial humanized nanobody libraries to identify compatible binders helps to counter the immunogenic defenses developed by viral pathogens. The method described in this study could also be expanded to infectious diseases other than COVID-19 and serve as a technical reserve for rapid neutralizing nanobodies discovery during future pandemics.

## Acknowledgments

We thank all colleagues of the COVID-19 team from NCATS for their support during the study, including Ganesha Bantukallu and Vinoth Chenniappan for help with lyophilizations Bolormaa Baljinnyam for help with initial BLI and nano-DSF efforts, and Paul Shinn for his advice and guidance. We thank Drs. Andrew C. Kruse (Harvard University) and Aaron M. Ring (Yale University) for support and suggestions for constructing the phage libraries. We thank Vanessa Wall, Kelly Snead, Jennifer Mehalko, Matthew Drew, and Simon Messing of the Frederick National Laboratory for scientific support of protein production efforts. We thank Dr. Gary Whittaker (Cornell University) for providing plasmids for pseudotyping.

## Supplementary Materials

Fig S1. Nanobody purification and quality control

Fig S2. Bio-layer interferometry binding profiles for immobilized nanobodies binding RBD-mFc

Fig S3. Bio-layer interferometry binding profile for immobilized nanobodies binding S1-hFc

Fig S4. QD ACE2-GFP endocytosis assay with nanobody treatment

Fig S5. QD ACE2-GFP endocytosis assay with Fc treatment

Fig S6. Global fit curves of 1-2G-Fc and 2-1F-Fc binding to RBD-mFc

Fig S7. SARS-CoV-2 pseudotyped particles assay for non-blocks and the trimer

Fig S8. Cross-reactivity screen of RBD-1-2G

Fig S9. Effect of lyophilization on RBD-1-2G

Fig S10. Atomic fit models of the RBD/ACE2/Nanobody interactions

Fig S11. CryoEM workflow chart and refinement strategy

Fig S12. Root mean square deviation (RMSD) of the MD triplicates

Fig S13. Root mean square fluctuation (RMSF)

Fig S14. Hydrogen bonds and salt bridge interactions per time for the MD stimulations

Fig S15. Free energy of RBD-1-2G binding

Fig S16. Cryo-EM structure of RBD-1-2G bound to B.1.1.7 S protein.

Table S1. Nanobody complementarity-determining regions (CDRs) sequences and phage enrichment stats.

Table S2. Microscope parameters used to collect Cryo-EM data of the nanobody complexes

Table S3. Hydrogen bond distances.

## Author contributions

Conceptualization: YF, MH, MJB, AS

Methodology: YF, BDF, MH, MJB, AS

Investigation: YF, JFR, BDF, AR, CZC, XH, KG, QH, EML, RP, MX, MP, RE, ZI, TS, AH, VD, WG, TT, NR, SP, DE, EO, KS, MW

Visualization: BDF, YF, JFR, AR, MJB

Writing – original draft: YF, BDF, JFR, AR, MH, MJB, AS

Writing – review & editing: BDF, YF, JFR, AR, MH, MJB, AS

Funding acquisition: MH, MJB, AS, MF

Project administration: MH, MJB, AS

Supervision: YF, BDF, MF, MH, MJB, AS

## Data and materials availability

All data, code, and materials used in the analysis must be available in some form to any researcher for purposes of reproducing or extending the analysis. Include a note explaining any restrictions on materials, such as materials transfer agreements (MTAs). Note accession numbers to any data relating to the paper and deposited in a public database; include a brief description of the data set or model with the number. If all data are in the paper and supplementary materials, include the sentence “All data are available in the main text or the supplementary materials.”

