## Supplemental Material for "A humanized nanobody phage display library yields potent binders of SARS CoV-2 spike"

Ying Fu^1¶^, Juliana da Fonseca Rezende e Mello^2¶^, Bryan D. Fleming^1¶^, Alex Renn^1¶^, Catherine Z. Chen^1^, Xin Hu^1^, Miao Xu^1^, Kirill Gorshkov^1^, Quinlin Hanson^1^, Wei Zheng^1^, Emily M. Lee^1^, Lalith Perera^2^, Robert Petrovich^2^, Manisha Pradhan^1^, Richard T. Eastman^1^, Zina Itkin^1^, Thomas Stanley^2^, Allen Hsu^2^, Venkata Dandey^2^, William Gillette^3^, Troy Taylor^3^, Nitya Ramakrishnan^3^, Shelley Perkins^3^, Dominic Esposito^3^, Eunkeu Oh^4^, Kimihiro Susumu^5^, Mason Wolak^4^, Marc Ferrer^1^_,_ Matthew D. Hall^1^*, Mario J. Borgnia^2^*, and Anton Simeonov^1^*

^1^ National Center for Advancing Translational Sciences, National Institutes of Health, Rockville, Maryland, USA.

^2^ Genome Integrity and Structural Biology Laboratory, National Institute of Environmental Health Sciences, National Institutes of Health, Department of Health and Human Services, Research Triangle Park, NC, USA.

^3^ Protein Expression Laboratory, NCI RAS Initiative, Cancer Research Technology Program, Frederick National Laboratory for Cancer Research, Frederick, MD, USA.

^4^ Optical Sciences Division, Code 5600, Naval Research Laboratory, Washington, D.C., USA.

^5^ Jacobs Corporation, Hanover, Maryland, USA.

^¶^These authors contributed equally: Ying Fu, Juliana da Fonseca Rezende e Mello, Bryan D Fleming and Alex Renn

**Supplemental Material contents:**

Figure S1-S16

Table S1-S3

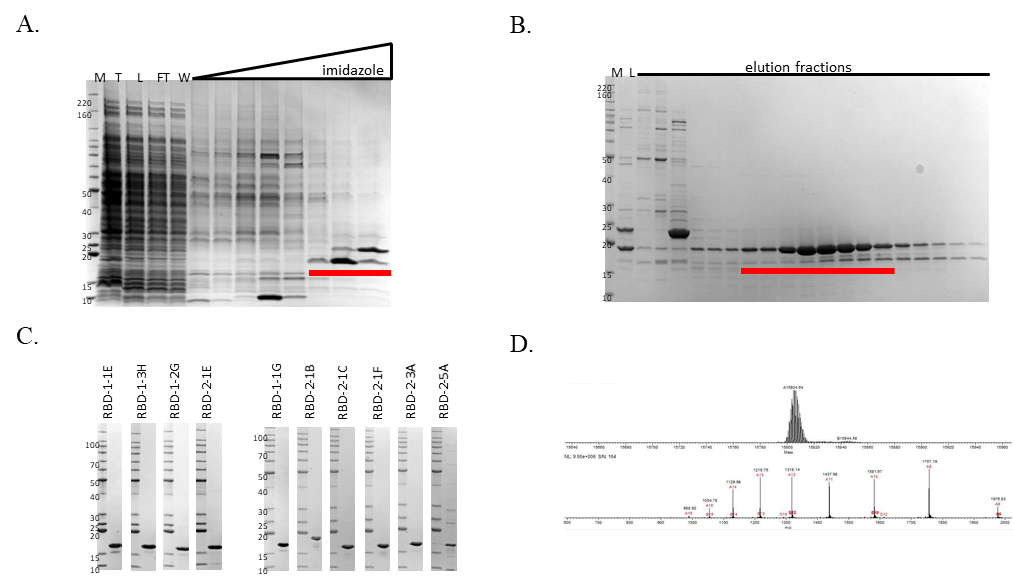

Figure S1: Nanobody purification and quality control. (A) Representative IMAC purification. T – total protein (lysate), L – column load (clarified lysate), FT – column flow through, W – column wash (B) Representative preparative SDS-PAGE after size exclusion column purification (SEC). L – column load. (C) SDS-PAGE Coomassie Blue staining of purified nanobodies. (D) Electrospray ionization mass spectrometry (ESI-MS) of a representative purified nanobody. All gels in panels A-C are SDS-PAGE/Coomassie staining with mass of protein standards noted in kDa. Red bars indicate fractions pooled.

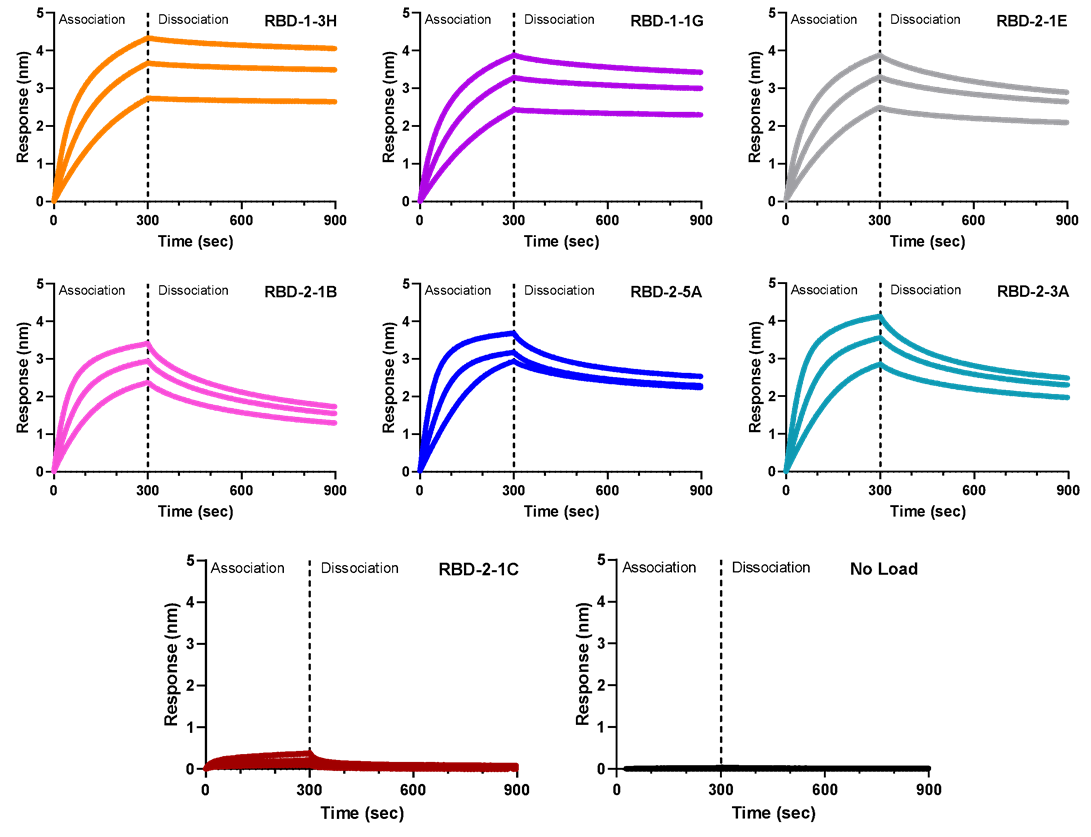

Figure S2: Octet Binding Profiles for immobilized nanobodies binding RBD-mFC for 200 nM, 100 nM and 50 nM concentrations.

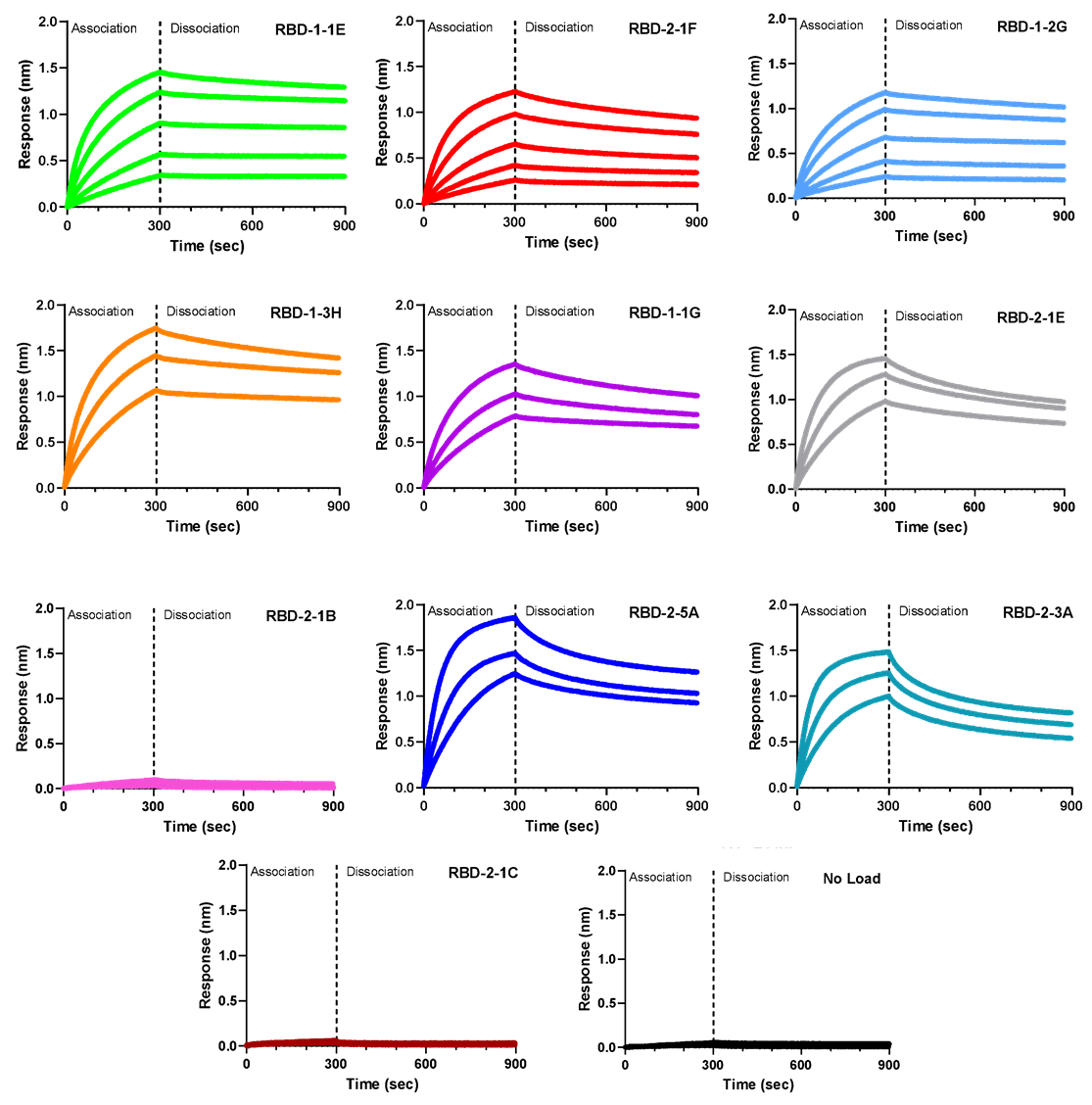

Figure S3: Octet Binding Profiles for immobilized nanobodies binding S1-hFc for 200 nM, 100 nM and 50 nM. RBD-1-2G, RBD-2-1F and RBD-1-1E also have 25 nM and 12.5 nM conditions.

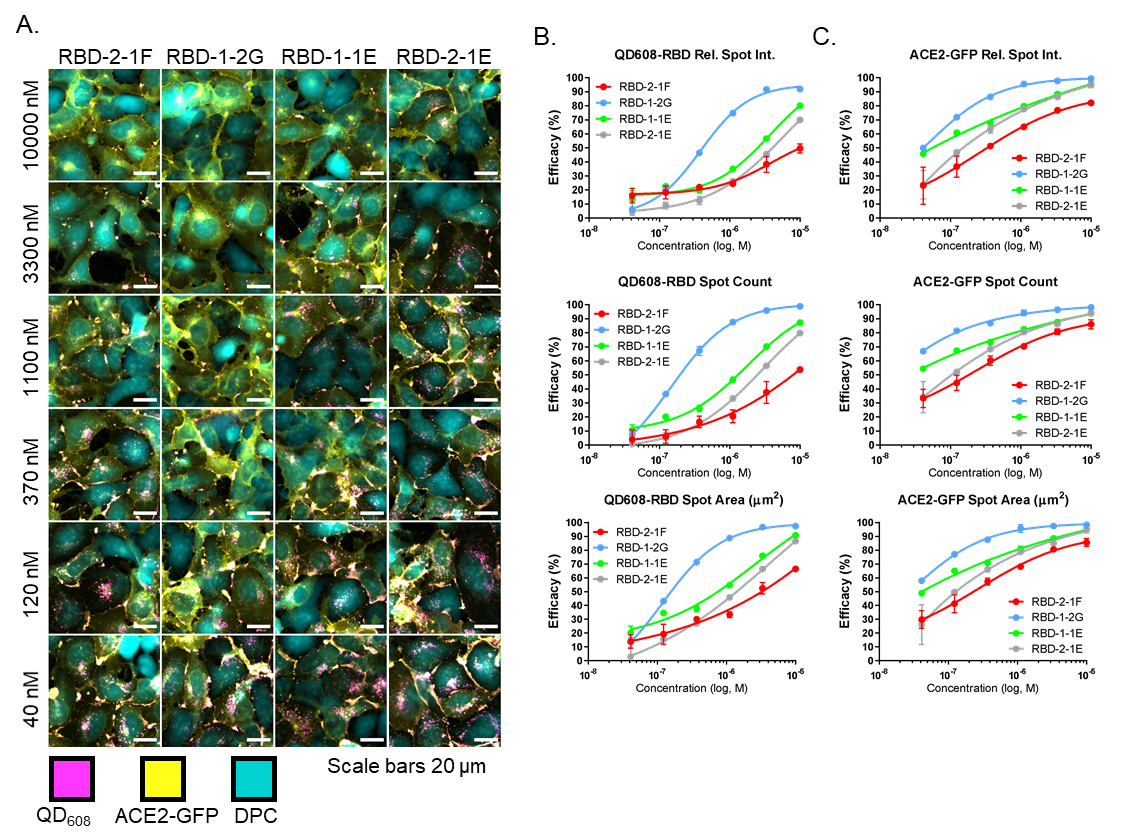

Figure S4: QD ACE2-GFP endocytosis assay with nanobody treatment. (A) Representative images for nanobody inhibition of QD endocytosis. Representative image montage of ACE2-GFP (yellow) HEK293T cells treated with QD_608_-RBD (magenta) that was preincubated for 30 minutes with RBD-2-1F, RBD-1-2G, RBD-1-1E, and RBD-2-1E starting at a concentration of 10 µM. Cells were treated for a total of three hours. Digital Phase Contrast (DPC, cyan) was used to visualize cell bodies. Scale bar, 20 µm. (B-C) Quantification of (B) QD_608_-RBD and (C) ACE2-GFP using high-content image analysis in each channel. Data was normalized to Optimem I treated cells (100%) and QD608-RBD alone (0%). N= approximately 2500 cells from duplicate wells, representative of three independent experiments. Curves fit using non-linear regression. Error bars indicate S.D.

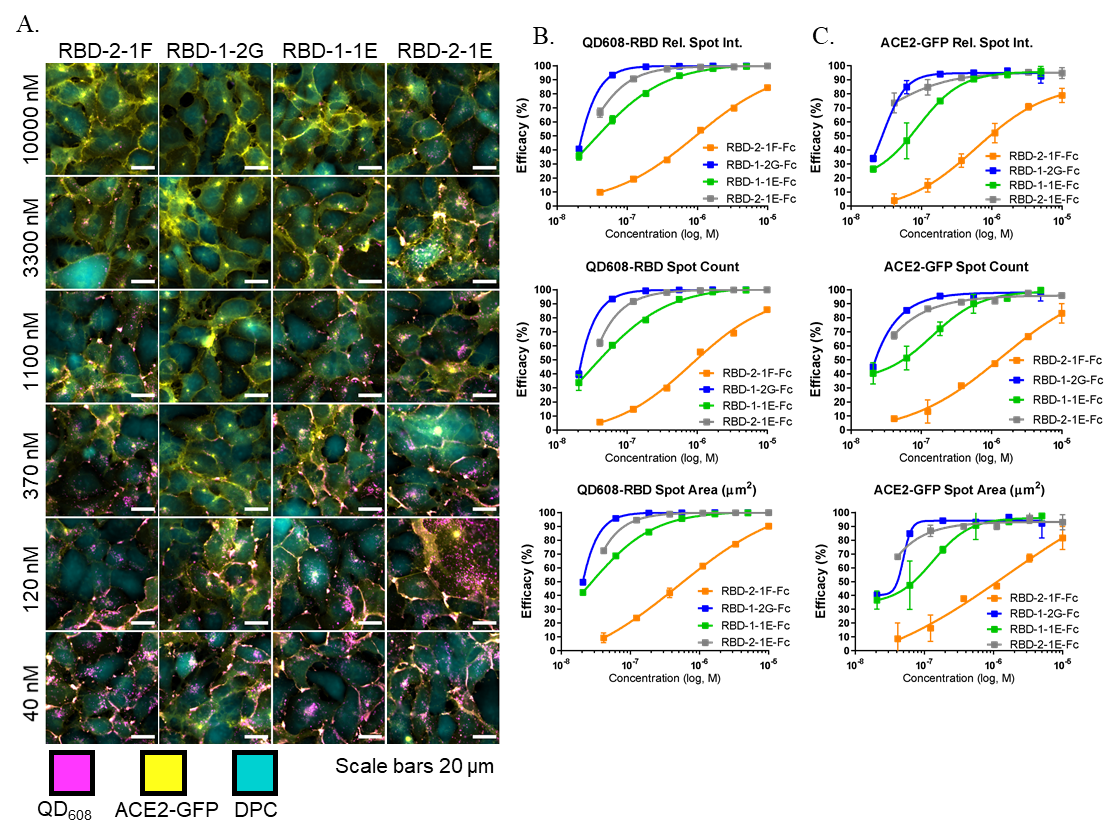

Figure S5: QD ACE2-GFP endocytosis assay with Fc treatment. (A) Representative images for Fc inhibition of QD endoytosis. Representative image montage of ACE2-GFP (yellow) HEK293T cells treated with QD_608_-RBD (magenta) that was preincubated for 30 minutes with RBD-2-1F-Fc, RBD-1-2G-Fc, RBD-1-1E-Fc, and RBD-2-1E-Fc starting at a concentration of 5 µM or 10 µM. Cells were treated for a total of three hours. Digital Phase Contrast (DPC, cyan) was used to visualize cell bodies. Scale bar, 20 µm. (B-C) Quantification of (B) QD_608_-RBD and (C) ACE2-GFP using high-content image analysis in each channel. Data was normalized to Optimem I treated cells (100%) and QD608-RBD alone (0%). N= approximately 1600 cells from duplicate wells, representative of three independent experiments. Curves fit using non-linear regression. Error bars indicate S.D.

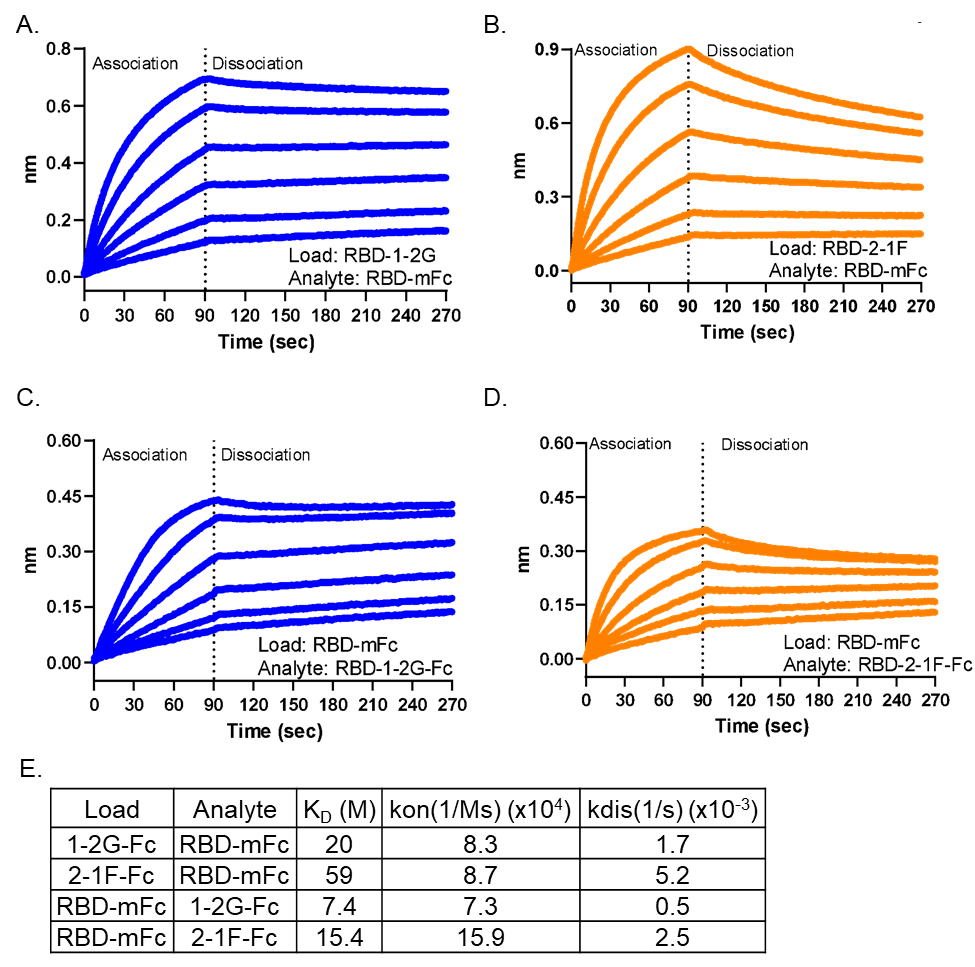

Figure S6: Global Fit curves of 1-2G-Fc and 2-1F-Fc binding to RBD-mFc. (A-B) Octet binding profiles using either RBD-1-2G-Fc(A) or RBD-2-1F-Fc(B) as load protein. Association of RBD-mFc ranging from 200 nM to 6.25 nM (1:2 serial dilutions) were used for global curve fitting. (C-D) Octet biosensors were loaded with RBD-mFc, then exposed to various concentrations of RBD-1-2G-Fc(C) or RBD-2-1F-Fc(D)(200 nM to 6.25 nM, 1:2 serial dilutions). (E) Calculations from global fit modeling for panels A-D.

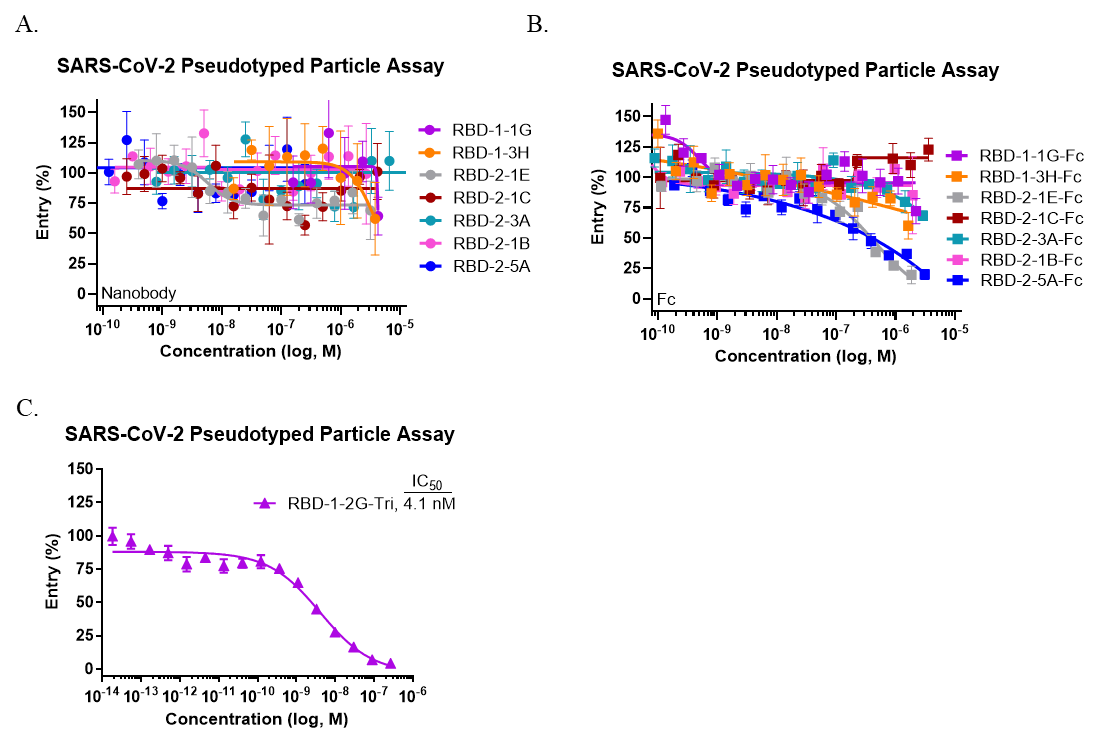
Figure S7: SARS-CoV-2 Pseudotyped particles assay for non-blockers and the trimer. SARS-CoV-2 pseudotyped particle entry assay for Nanobody(A), Fc(B) and Trimer(C) formats.

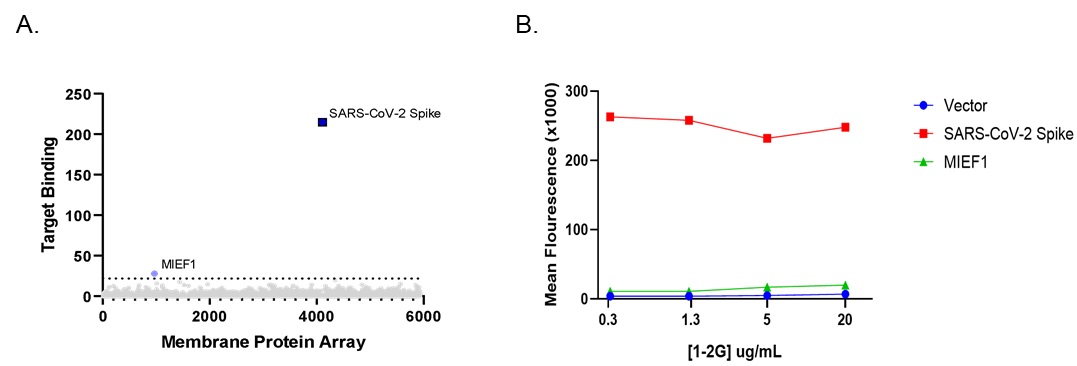

Figure S8: (A-B) Cross-reactivity screen of RBD-1-2G. (A) Human membrane proteome array (MPA) results identified during a cross-reactivity screen. (B) Follow-up validation screen to determine the reactivity of MIEF1 protein to RBD-1-2G.

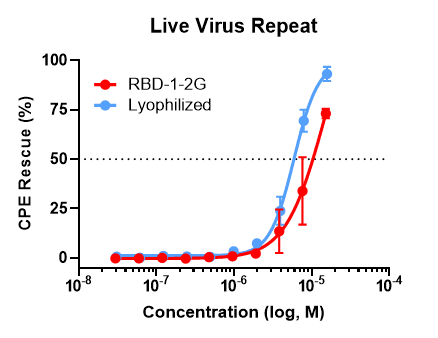

Figure S9: Effect of lyophilization on RBD-1-2G. RBD-1-2G vs a lyophilized reconstituted sample were tested for their ability to inhibit live virus infection. Titration curves with a top concentration of 15.2 µM for the untreated RBD-1-2G and 15.8 µM for the reconstituted lyophilized RBD-1-2G sample. Samples were diluted 1:2 in dPBS before being used for the viral assay. Technical replicates are n = 3 per concentration, all error bars represent S.D.

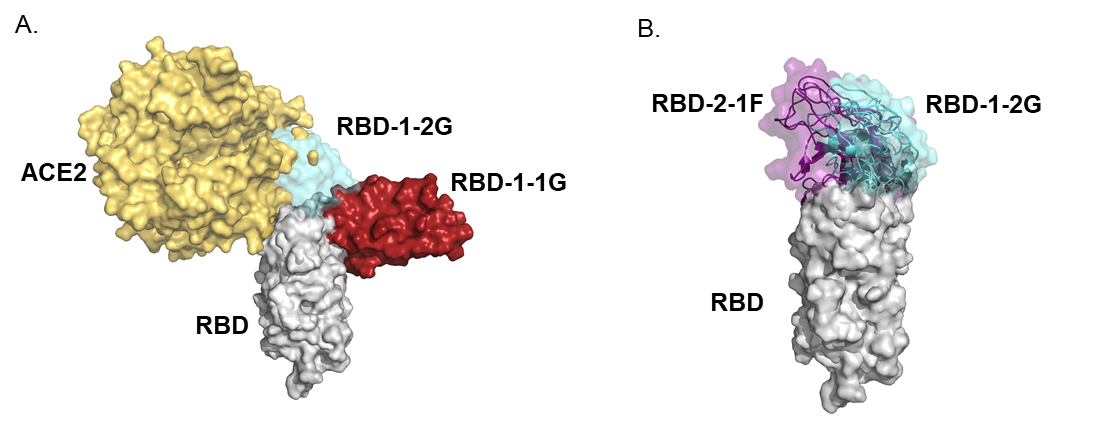

Figure S10 : Atomic Fit Models of the RBD/ACE2/Nanobody interactions. (A) Model of RBD-1-2G binding overlaps with the ACE2 binding site, while RBD-1-1G fails to inhibit binding. (B) Overlap of RBD-2-1F and RBD-1-2G suggesting similar epitopes being targeted by ‘Group 1’ binders.

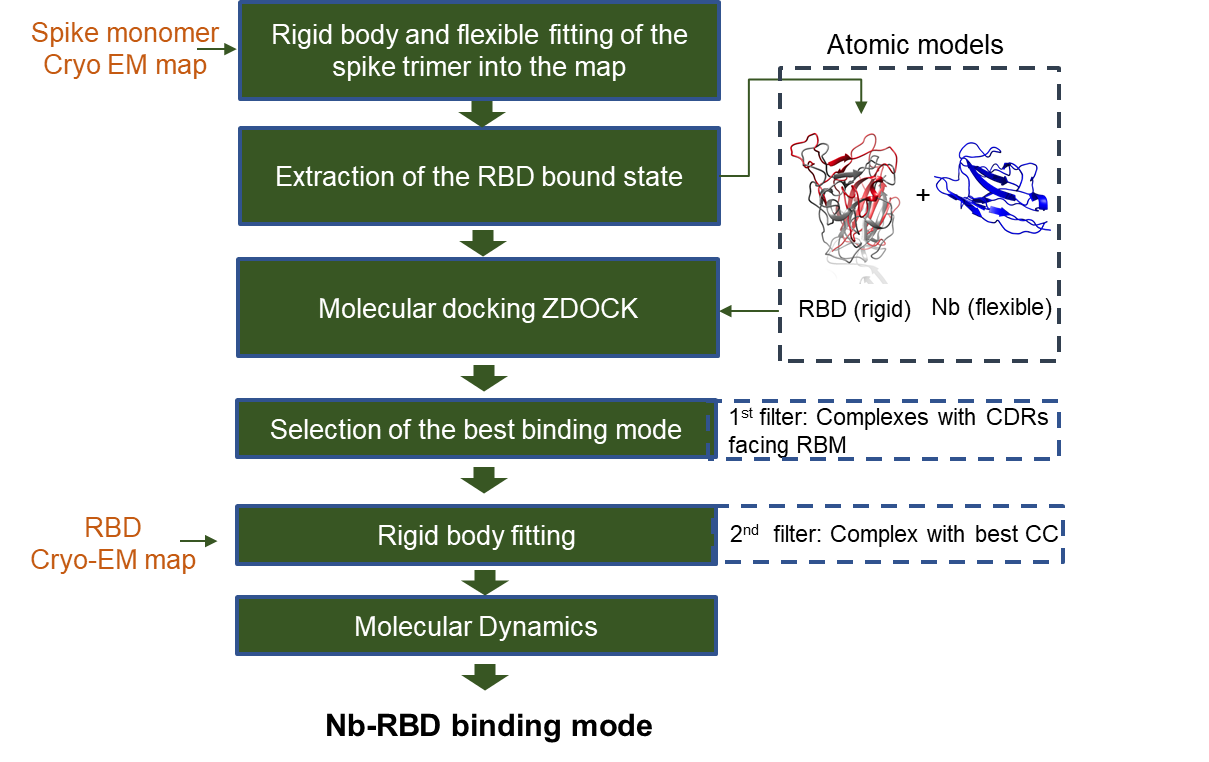

Figure S11: Workflow describing the combination of computational techniques and Cryo-EM applied to identify Nb-RBD binding mode details.

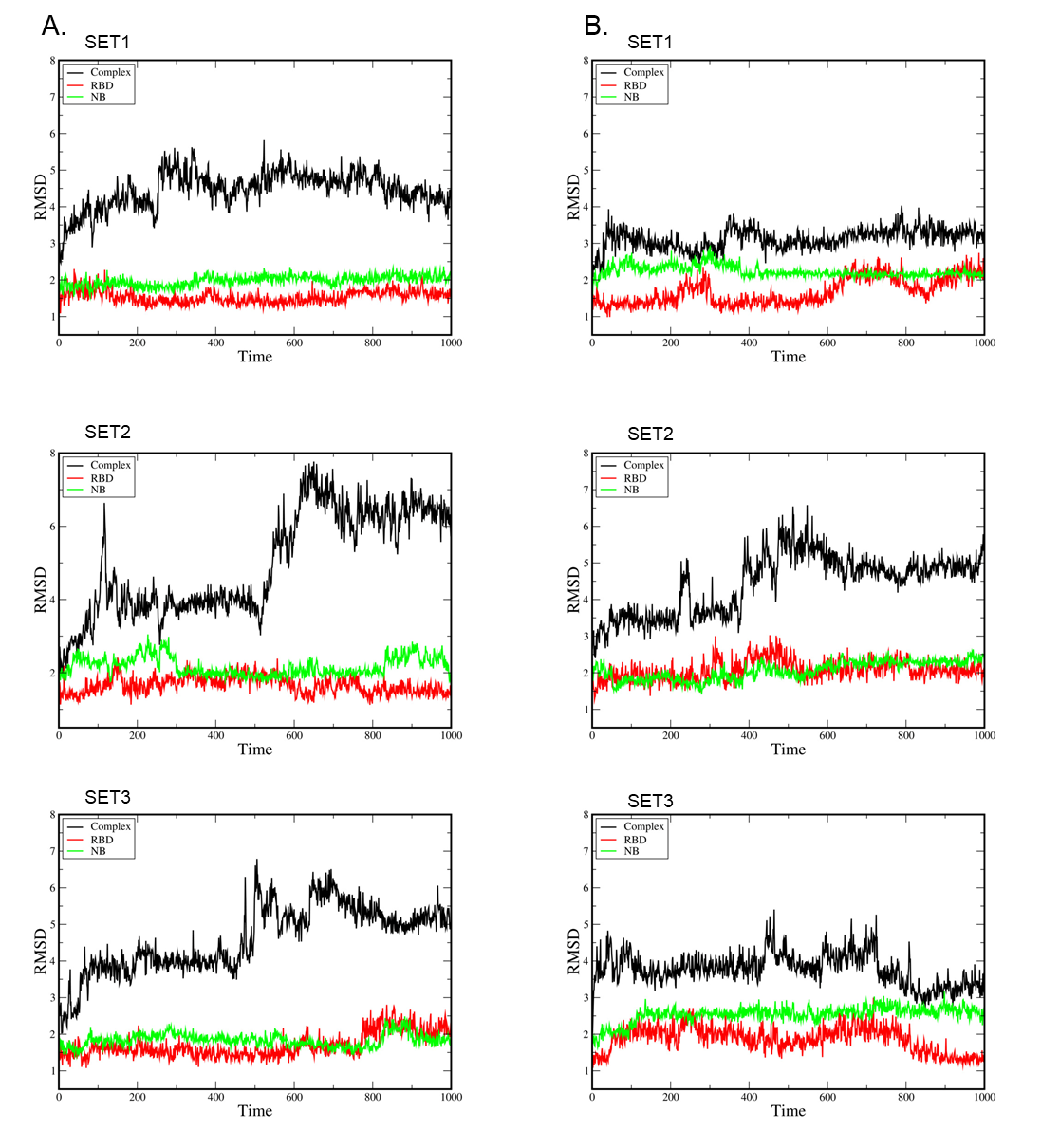

Figure S12: Root mean square deviation (RMSD) of the MD triplicates per column for RBD-1-2G in complex (A) with the WT RBD and (B) with B.1.1.7 RBD showed that the systems were stable.

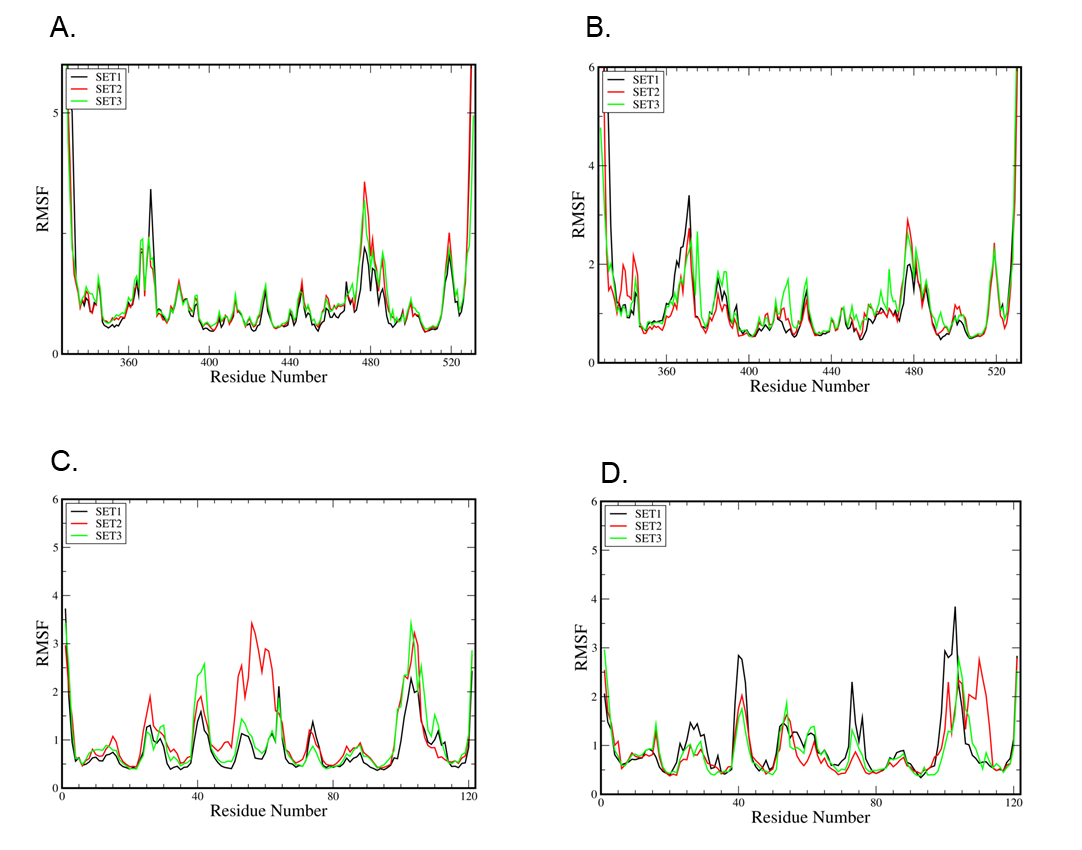

Figure S13: Root mean square fluctuation (RMSF) for the (A) WT RBD residues and (B) B.1.1.7 RBD residues, showing the same pattern of variations. Both graphs show two main peaks correspondent to the residues 355-375 and 470-490. (C) The fluctuation of the residues of the RBD-1-2G in complex with the WT RBD and (D) with the B.1.1.7 RBD.

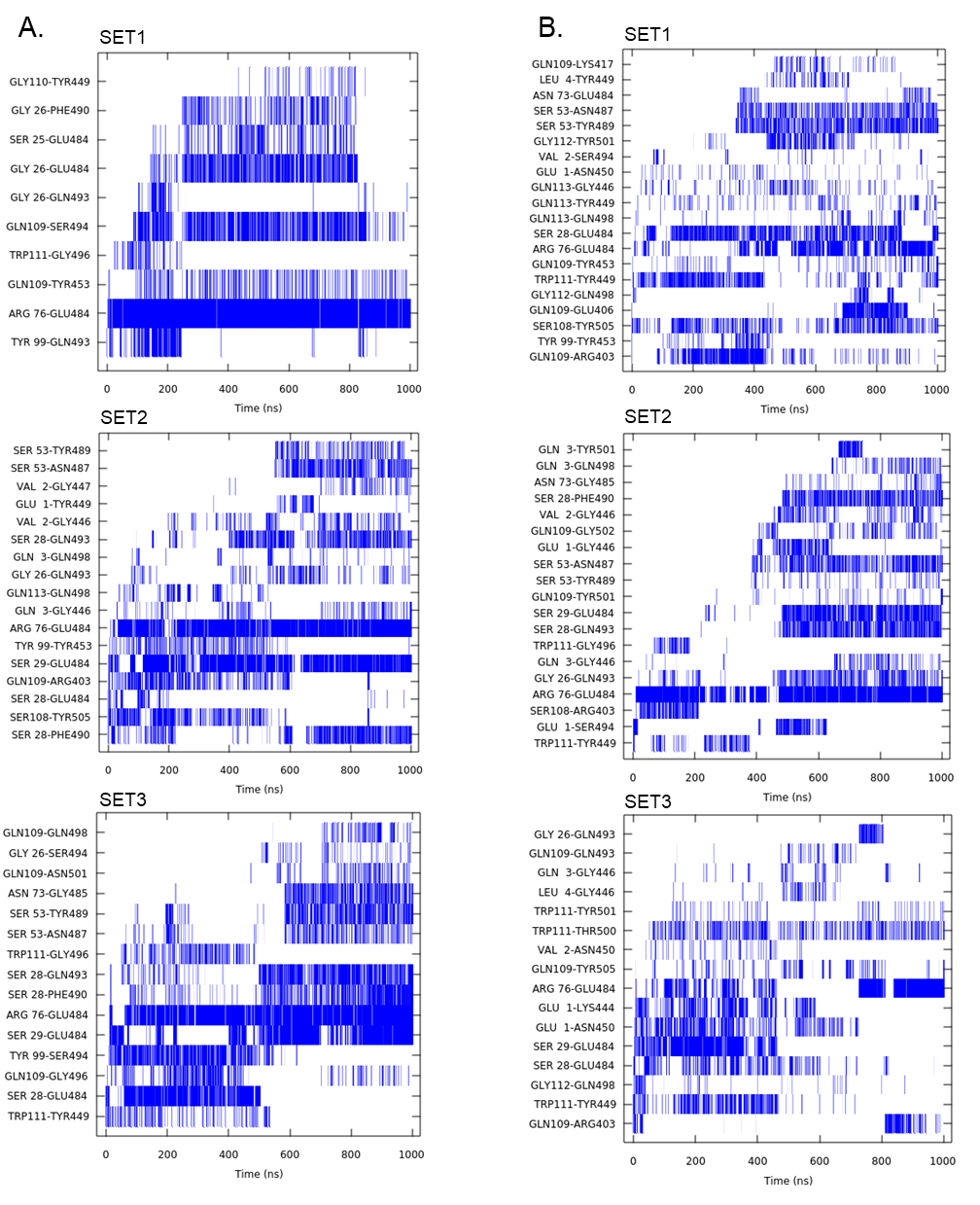

Figure S14: Hydrogen bonds and salt bridge interactions per time of the MD simulations in triplicate for the systems (per column) containing RBD-1-2G with (A) WT RBD and (B) with B.1.1.7 RBD.

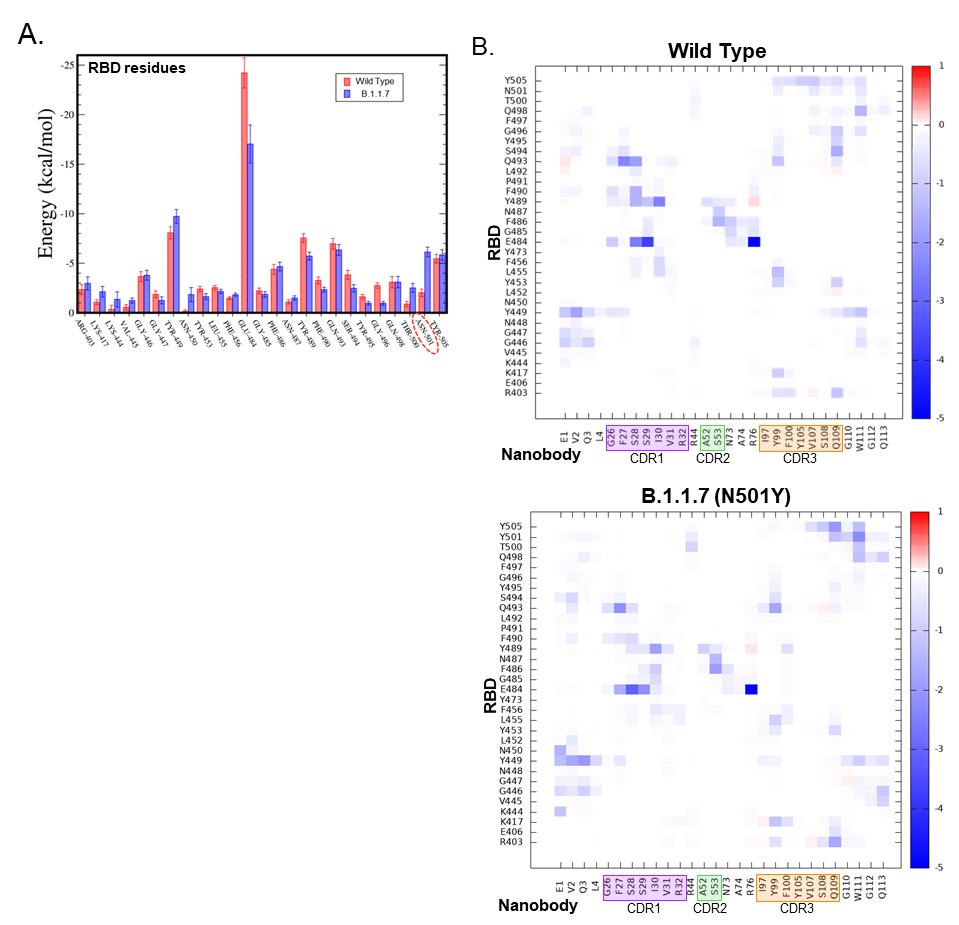

Figure S15: Free energy of binding for RBD-1-2G (A) Total free energy of binding each residue in the WT and B.1.17 variant RBD when bound by RBD-1-2G. (B) Heatmap showing the free binding energy for the WT and B.1.1.7 RBD in complex with RBD-1-2G.

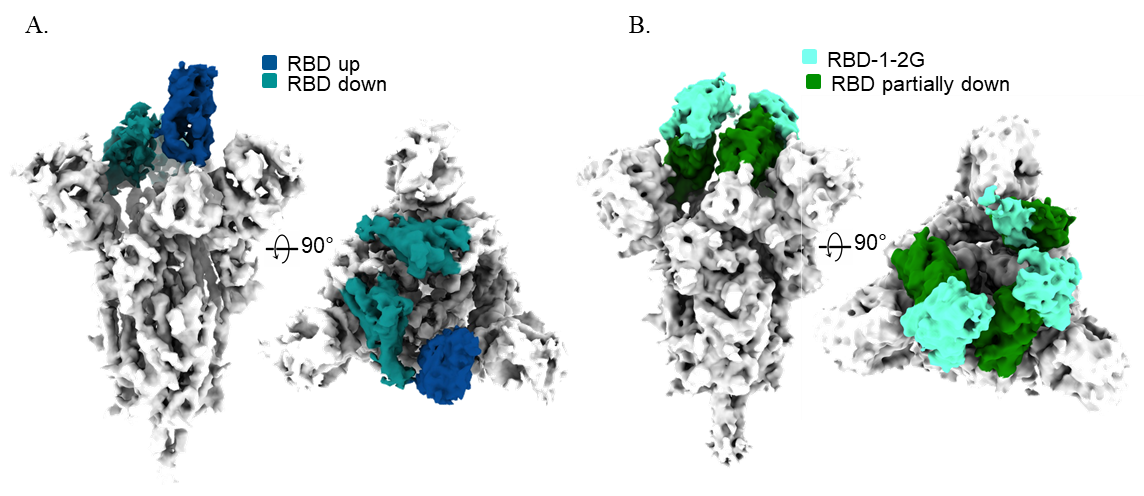

Figure S16: Cryo-EM structure of the (A) APO state WT SARS-CoV-2 S-protein in side and top views showing the RBD in the 1 Up, 2 down state. (B) B.1.1.7 SARS-CoV-2 S-protein bound to RBD-1-2G in side and top views showing the RBD in the partially down state.

| **Nanobody** | **CDR1 Region** | | **CDR2 Region** | | | **CDR3 Region** | **Sequence Enrichment** |
| --- | --- | --- | --- | --- | --- | --- | --- |
|  | Lead | Variable | Lead | Variable | Tail | Variable |  |
| RBD-1-3H | GNIS | ADAH | IT | NP | GGNTN | SLLFKRNELSNRWT | 8 / 48 (16.7%) |
| RBD-1-1E | GYIS | RSVF | IS | PD | GANTN | IIRYVASNNTEDV | 6 / 48 (12.5%) |
| RBD-1-2G | GFSS | IVY | ID | AS | GSTTN | IAYFTSPEYVVSQG | 6 / 48 (12.5%) |
| RBD-1-1G | GSIS | EYKH | IT | NP | GSNTN | VLFPKTATKNVGWE | 6 / 48 (12.5%) |
| RBD-1-2E | GNIS | AHGV | ID | RP | GANTN | GYTWVAQTQHYSWE | 1 / 48 (2.1%) |
| RBD-2-1B | GYIS | VTGI | IT | VY | GTNTY | KLKSDPVNEPSPEE | 26 / 95 (27.4%) |
| RBD-2-1F | GYIS | Q | ID | QD | GTSTN | ANFLVPGQLRKI | 25 / 95 (26.3%) |
| RBD-2-5A | GNIS | RIVY | IA | AQ | GTSTN | LHTYVAAEKIVEQH | 8 / 95 (8.4%) |
| RBD-2-1C | GYIS | SVSS | IT | AL | GTSTY | GWDFTNAKGVIHET | 7 / 95 (7.4%) |
| RBD-2-1E | GTIS | HETF | ID | AS | GTNTN | VAYFPVLTTQVISL | 5 / 95 (5.3%) |
| RBD-2-3A | GSIS | QAFI | IS | TE | GSSTN | LRKFQPDIQLTQWV | 4 / 95 (4.2%) |
| RBD-2-3H | GSIS | SHEQ | IT | FI | GGNTY | ALIREDSLSKTHWE | 2 / 95 (2.1%) |
| RBD-2-2C | GTIF | HEVW | ID | VI | GTSTY | SGVHQGRRHVKFLV | 1 / 95 (1.1%) |

Supplemental Table 1: Nanobody complementarity-determining regions (CDRs) sequences and phage enrichment stats

| **Data set** | **Voltage (KeV)** | **Mag (kX)** | **Pixel Size (Å)** | **Underfocus Range (Å)** | **Exposure Rate (e/px/s)** | **Total Dose** | **Movies Collected** |
| --- | --- | --- | --- | --- | --- | --- | --- |
| **RBD-1-2G** | 300 | 130 | 0.532 (SR) | 1.2 – 2.4 | 8 | 60 | 4230 |
| **RBD-1-1G** | 200 | 36 | 1.187 (CM) | 1.2 – 2.2 | 8 | 60 | 2743 |
| **RBD-1-3H** | 200 | 36 | 1.187 (CM) | 1.5 – 2.8 | 8 | 60 | 4977 |
| **RBD-2-1F** | 200 | 36 | 1.187 (CM) | 1.2 – 2.8 | 8 | 60 | 3347 |
| **RBD-2-1B** | 200 | 36 | 1.187 (CM) | 1.4 – 2.4 | 8 | 60 | 1693 |
| **RBD-2-3A** | 200 | 36 | 1.187 (CM) | 1.5 – 2.2 | 8 | 60 | 1755 |

Supplemental Table 2: Microscope parameters used to collect Cryo-EM data of the nanobody complexes.

| **Complex** | **Interactions** | **Distance (Å)** |
| --- | --- | --- |
| RBD (WT)  + RBD-1-2G | E484-R76 (1) | 2.8 |
|  | E484-R76 (2) | 2.6 |
|  | E484-S25 | 2.7 |
|  | Y489-S29 | 3.3 |
|  | N501-Y99 | 3.1 |
|  | N501-Q498 | 3.2 |
|  | N501-G496 | 3.0 |
| RBD (B.1.1.7)  + RBD-1-2G | E484-R76 (1) | 3.0 |
|  | E484-R76 (2) | 2.7 |
|  | E484-S28 | 2.8 |
|  | W111-T500 | 3.3 |
|  | Y501-G112 | 2.7 |
|  | Y501-Q498 | 3.2 |

Supplemental Table 3: Distance of most relevant intra- and inter-molecular hydrogen bonds in complexes containing RBD-1-2-G with RBD (WT) or RBD (B.1.1.7).
